# Molecular Diversity Among Adult Human Hippocampal and Entorhinal Cells

**DOI:** 10.1101/2019.12.31.889139

**Authors:** Daniel Franjic, Jinmyung Choi, Mario Skarica, Chuan Xu, Qian Li, Shaojie Ma, Andrew T. N. Tebbenkamp, Gabriel Santpere, Jon I. Arellano, Ivan Gudelj, Lucija Jankovic-Rapan, Andre M. M. Sousa, Pasko Rakic, Nenad Sestan

## Abstract

The hippocampal-entorhinal system is comprised of functionally distinct subregions collectively critical for cognition, and selectively vulnerable to aging and pathological processes. To gain insights into neuronal and non-neuronal populations within this system, we performed single-nucleus transcriptional profiling from five human hippocampal-entorhinal subregions. We found that transcriptomic diversity of excitatory neurons across these subregions reflected the molecular transition from three-layered archicortex to six-layered neocortex. Additionally, mRNA and protein for DCX, an immature neuron marker, were clearly detected in some cells, but not in dentate granule cells, the cell-type predicted to be generated in adult neurogenesis. We also found that previously functionally uncharacterized METTL7B was enriched in human and non-human primate neuronal subtypes less vulnerable to initial Alzheimer’s disease pathology. Proteomic and biochemical assays revealed METTL7B interacts with Alzheimer’s disease-related proteins, including APP, and its overexpression reduced amyloid-beta generation. These results reveal cell type-specific molecular properties relevant for hippocampal-entorhinal physiology and dysfunction.

## INTRODUCTION

The neural circuits of the hippocampal formation (HIP) and entorhinal cortex (EC) are critical components of a widespread neural network for memory and representation of space and time (Gloor, 1997; Andersen, 2007; Buzsaki and Moser, 2013). Based on cytoarchitectonic, cellular, and circuitry variations, the hippocampal-entorhinal system can be subdivided into functionally distinct subregions that gradually transition from the simple three-layered dentate gyrus (DG) and hippocampus (Cornu Ammonis, CA), and through more complex lamination of the subiculum (collectively referred to as the allocortex) to the six-layered EC (mesocortex) (Freund, 2002; Suzuki and Amaral, 2004; Klausberger and Somogyi, 2008). The molecular basis of the diversity of cell types in these subregions and their homology with bordering neocortical cell types and lamination remains poorly understood (Kriegstein and Connors, 1986; Hoogland and Vermeulen-Vanderzee, 1989; Reiner, 1991; Ishizuka, 2001; Zeisel et al., 2015; Cembrowski et al., 2016b; Mercer and Thomson, 2017; Shepherd and Rowe, 2017).

Primate- or human-specific evolutionary innovations may underlie some region-selective aspects of hippocampal cell types, necessitating the study specifically of the human hippocampus. The cytoarchitectonic differentiation of the allocortex and neocortex arose very early during mammalian evolution, with the mammalian allocortex reminiscent of the three-layered reptilian cortex rather than six-layered mammalian cortex. Based on this similarity, as well as histological, physiological and connectional studies, it is hypothesized that allocortex is composed of excitatory projection neurons that resemble those specifically in the deep layers of the mammalian neocortex (Kriegstein and Connors, 1986; Reiner, 1991; Ishizuka, 2001; Luzzati, 2015; Shepherd and Rowe, 2017). However, although the hippocampus is a phylogenetically ancient part of the cerebral cortex, the hippocampal-entorhinal system and its cortico-cortical projections are greatly expanded in humans and non-human primates (Stephan, 1975; Demeter et al., 1990; Morrison and Hof, 1997; Suzuki and Amaral, 2004; Patzke et al., 2015). Moreover, the human and non-human primate homologs of this system have undergone extensive evolutionary changes also in gene expression and perhaps cell composition as compared to some commonly studied mammals such as rodents (Bakken et al., 2015; Sousa et al., 2017). As a potential consequence, the normal functions and disease features of cell types in these subregions may be absent or exhibit substantial differences in other species. For example, a distinctive feature of the hippocampal-entorhinal systems of most analyzed mammals (apart from cetaceans) is persistent adult neurogenesis of excitatory granule neurons in the DG (Patzke et al., 2015). However, whether new granule cells are generated in the adult human DG, and whether these neurons express DCX, a marker for immature neurons that is associated with neurogenesis, has not been fully resolved (Eriksson et al., 1998; Rakic, 2002; Spalding et al., 2013; Boldrini et al., 2018; Kempermann et al., 2018; Sorrells et al., 2018; Moreno-Jimenez et al., 2019).

Within the human hippocampal-entorhinal system, some cell types and circuits are selectively vulnerable in normal aging and certain pathological processes. For example, excitatory projection (pyramidal) neurons in the hippocampal CA1 field (Sommer’s sector), and to a lesser degree those in CA2-4 fields, are more vulnerable to hypoxia-ischemia damage and intractable seizures (mesial temporal sclerosis) than DG granule cells or other major hippocampal neuronal subtypes (Schmidt-Kastner and Freund, 1991; Blumcke et al., 2007). Alzheimer disease’s pathology, which is defined by extracellular accumulation of Aβ peptides and intracellular aggregates of hyperphosphorylated tau protein, also exhibits regional and cellular differences (Fischer, 1907; Glenner and Wong, 1984; Alzheimer et al., 1995; Hardy and Selkoe, 2002; Tanzi and Bertram, 2005; Ballatore et al., 2007; Karran and De Strooper, 2016). In mesial temporal regions, pathology first appears in layer 2 of the EC and hippocampal CA1 field (McMenemey, 1940; Wilcock and Esiri, 1982; Morrison and Hof, 1997; Serrano-Pozo et al., 2011; Braak and Del Trecidi, 2015), while DG cells, CA2-4 pyramidal neurons and layer 5B pyramidal neurons in the EC (Davies et al., 1992; Jin et al., 2004; Schonheit et al., 2004; West et al., 2004; Ohm, 2007) are more resilient in the early stages of the disease. Given this selectivity, a more detailed molecular profiling of this system, will aid our understanding of human brain development and neuropsychiatric disease.

To gain new insights into cell populations and cell type-specific differences in gene expression, evolution, neurogenic capability, and variable disease susceptibility, we performed high-coverage single-nucleus RNA sequencing (snRNA-seq) on five anatomically defined subregions of the hippocampal-entorhinal system. These efforts, like similar recent efforts to transcriptomically characterize the postmortem adult human brain (Krishnaswami et al., 2016; Lake et al., 2016; Habib et al., 2017; Lake et al., 2018; Li et al., 2018; Hodge et al., 2019; Mathys et al., 2019; Schirmer et al., 2019; Velmeshev et al., 2019) (including pioneering profiling of HIP (Habib et al., 2017), identified a highly diverse set of neuronal and non-neuronal cell types with clear regional distinctions and implications for human brain function, evolution, and disease.

## RESULTS

### Transcriptomic diversity of hippocampal and entorhinal cells

To survey the transcriptomic diversity and functional specification of the mesial temporal cortex, we used snRNA-seq to profile five subregions of the hippocampal-entorhinal system collected from fresh frozen postmortem brains of clinically unremarkable human donors. These specimens were selected from a larger pool of postmortem human brains based on the quality of isolated nuclei and RNA. Taking into consideration dramatic cytoarchitectonic variations, we microdissected the hippocampal formation (DG, CA2-4, CA1, and subiculum) and EC for a total of five subregions (Fig. 1A).

**Figure 1.**
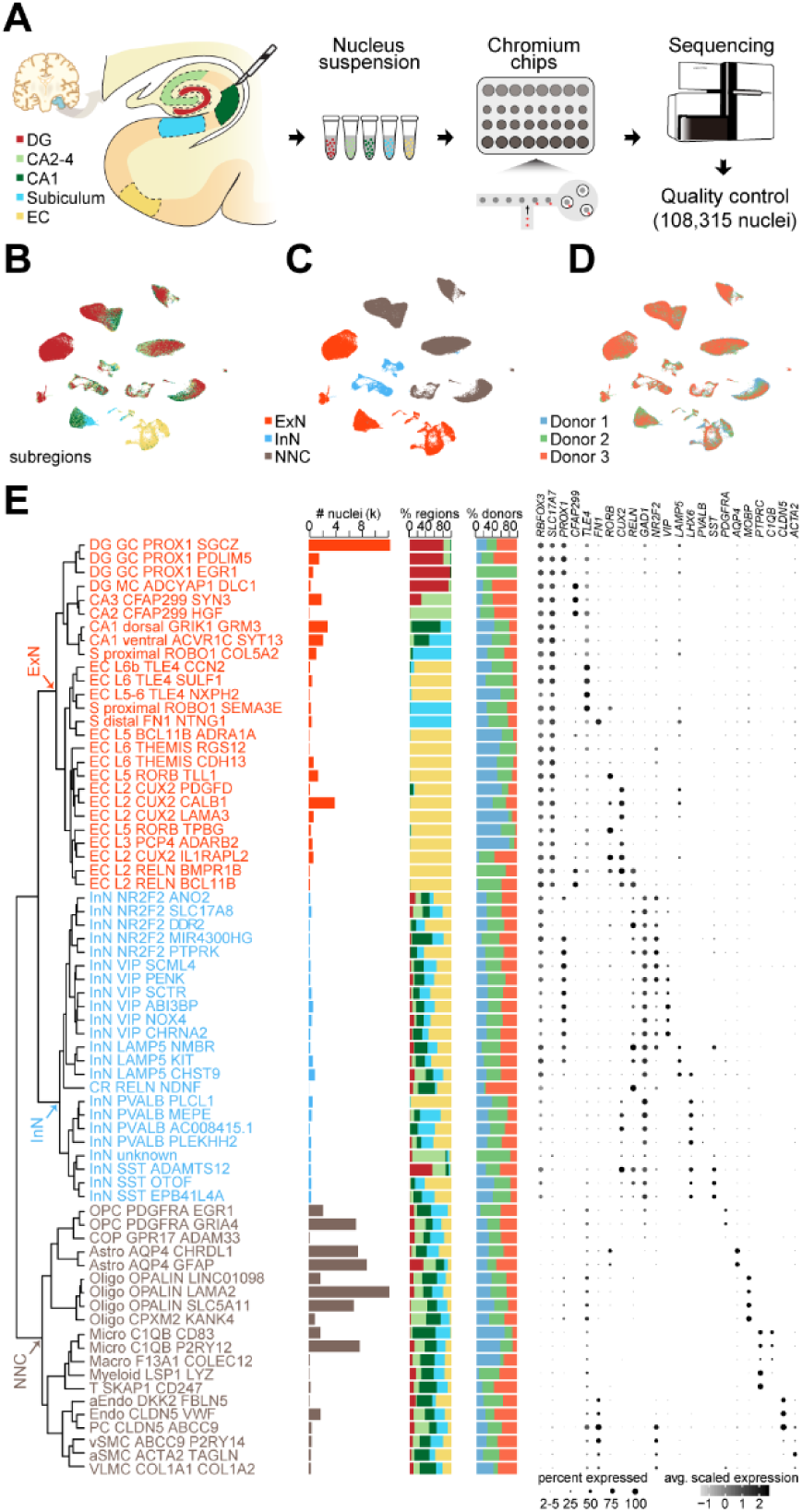
Cell type diversity in the hippocampal-entorhinal system revealed by single-nucleus RNA-seq. **A,** Scheme outlining the snRNA-seq procedure including subregion dissections of hippocampus and entorhinal cortex, nucleus isolation and capture, RNA sequencing and quality control. **B-D,** Uniform Manifold Approximation and Projection (UMAP) visualization representing the transcriptomic arrangement of all nuclei, colored by different subregions (B) major cell types (C), and donors (D). ExN, excitatory neurons; InN, inhibitory neurons; NNC, non-neuronal cells. **E.** Dendrogram depicting the hierarchical taxonomy across all cell subtypes. Bar plots show the number of nuclei, subregional and donor contributions within each subtype, with coloring scheme conforming to panel **c**. Dot plot demonstrates the expression of specific marker genes along the cell-type taxonomy. The size and color of dots indicate the percent of expressed nuclei and the average gene expression within each subtype, respectively. GC, granule cell; MC, mossy cell; OPC, oligodendrocyte precursor cell; COP, committed oligodendrocyte precursor cell; Astro, astrocyte; Oligo, oligodendrocyte; Micro, microglia; Macro, macrophage; Myeloid, myeloid cell; T, T cell; aEndo, arterial endothelial cell; Endo, endothelial cell; PC, pericyte; vSMC, venous smooth muscle cell; aSMC, arterial smooth muscle cell; VLMC, vascular and leptomeningeal cell. See also Figure S1 and S2.

Unbiased isolation of nuclei using our previously described protocol (Li et al., 2018; Zhu et al., 2018) followed by snRNA barcoding, cDNA sequencing and quality filtering yielded 108,315 high-quality single-nucleus profiles from all five subregions (Fig. 1A, S1A-D). Analysis of the expression of genes known to be enriched in major cell subpopulations suggested these included 44,697 neurons, of which 35,768 (80.02%) were glutamatergic excitatory neurons (expressing the gene encoding the vesicular glutamate transporter *SLC17A7*) and 8,929 (19.98%) were GABAergic inhibitory neurons (expressing the gene encoding the GABA synthesis enzyme *GAD1*), reflecting the expected 80:20 ratio of these populations. In addition, we identified 63,618 (58.73% of the total population) non-neuronal cells.

We next analyzed the transcriptomes of those nuclei on the Uniform Manifold Approximation and Projection (UMAP) layout representing their similarities at cellular granularity (Fig. 1B-D). Iterative clustering defined 69 transcriptomically distinct cell clusters representing presumptive cell types across all individuals (donors). These transcriptomically diverse subpopulations were organized into a dendrogramatic taxonomy reflecting their gene expression patterns and were subsequently assigned identities commensurate with predicted cell types. This allowed us to identify 26 subtypes of excitatory neurons (Fig. 1E **and** S2A-B), 23 inhibitory neuron subtypes (Fig. 1E **and** S2C-D), and 20 non-neuronal cell types and subtypes (Fig. 1E **and** S2E-F). Similar single nucleus approaches applied to human neocortical samples yielded comparable numbers and distributions of cell populations in medial temporal gyrus (MTG) (Hodge et al., 2019)and dorso-lateral prefrontal cortex (dlPFC) (Li et al., 2018)(Fig. S1E-F).

Within excitatory neuron subtypes, we found marked transcriptional diversity that reflects differences in the cytoarchitectonic organization among the subregions of the HIP (DG, CA2-4, CA1, and subiculum) and EC (Fig. 1E). For example, in addition to *ADCYAP1*-expressing mossy cells in DG, we found three distinct subclusters of *PROX1*-expressing granule cells. We also identified excitatory neurons in CA1 and CA2-4 that could be deconstructed into additional subtypes, indicating a finer molecular subdivision not readily apparent in the cytoarchitecture. Molecular distinctions were also evident in the subiculum, with two proximal subtypes close to CA1 expressing *ROBO1* and a third, distal subtype expressing *FN1* (Cembrowski et al., 2018). Within the EC, excitatory neurons were broadly characterized by laminar positioning. We identified seven neuron subtypes in layer 2/3 (characterized by high expression of *CUX2* and/or *RELN*) and eight subtypes in deep layers 5 and six with specific expression of deep-layer markers including *TLE4*, *ADRA1A*, and *THEMIS*.

In contrast to excitatory neurons, interneurons and non-neuronal cell types exhibited a more uniform spatial distribution (Fig. 1E). Transcriptomic diversity and multiple cell subtypes were evident among these populations, but this diversity did not generally segregate by percent region (middle histogram). For example, although the abundance of some interneuron subtypes differed between HIP and EC, all major subtypes of interneurons, including *SST*-, *PVALB*-, *VIP*-, and *LAMP5*-expressing interneurons, were shared across all subregions assayed. Among multiple non-neuronal cell types, we identified two astrocyte subtypes (Astro), two subtypes of oligodendrocyte precursor cells (OPCs), four subtypes of oligodendrocytes (Oligo), two subtypes of microglia (Micro), and multiple vasculature subtypes, each of which was generally shared across all of the five subregions dissected. These data therefore describe previously uncharacterized cell populations in the hippocampal-entorhinal system and extend previous findings concerning the functional specificity of neuronal and non-neuronal populations to this system (Freund, 2002; Suzuki and Amaral, 2004; Klausberger and Somogyi, 2008).

### Taxonomic relationships across allo-, meso- and neo-cortex

The putative homology between neurons in the hippocampal-entorhinal system and neocortical neurons, and in particular the cytoarchitectonic and evolutionary transition between allo-, meso-, and neo-cortex, offers an opportunity to reveal organizational principles underlying the specialization and function of the mammalian cerebral cortex. Towards elucidating these principles, we compared cell profiles across hippocampal-entorhinal subregions and transcriptomically defined cell types within two human neocortical regions (MTG and dlPFC).

Among hippocampal-entorhinal subregions, we observed a clear distinction between excitatory neurons of the hippocampus proper and DG, and the hippocampal formation more generally, as compared to those of EC (Fig. 2A-B, S2A-B). Excitatory neurons of the CA fields, subiculum and DG were also clearly distinct from those of MTG and dlPFC (Fig. 2B). We did, however, observe transcriptomic similarities between excitatory neurons in all neocortical layers in MTG and dlPFC (Fig. 3A). In contrast, we did not identify excitatory neurons in DG, CA fields, subiculum, and EC that corresponded to all neocortical layers in MTG and dlPFC.

**Figure 2.**
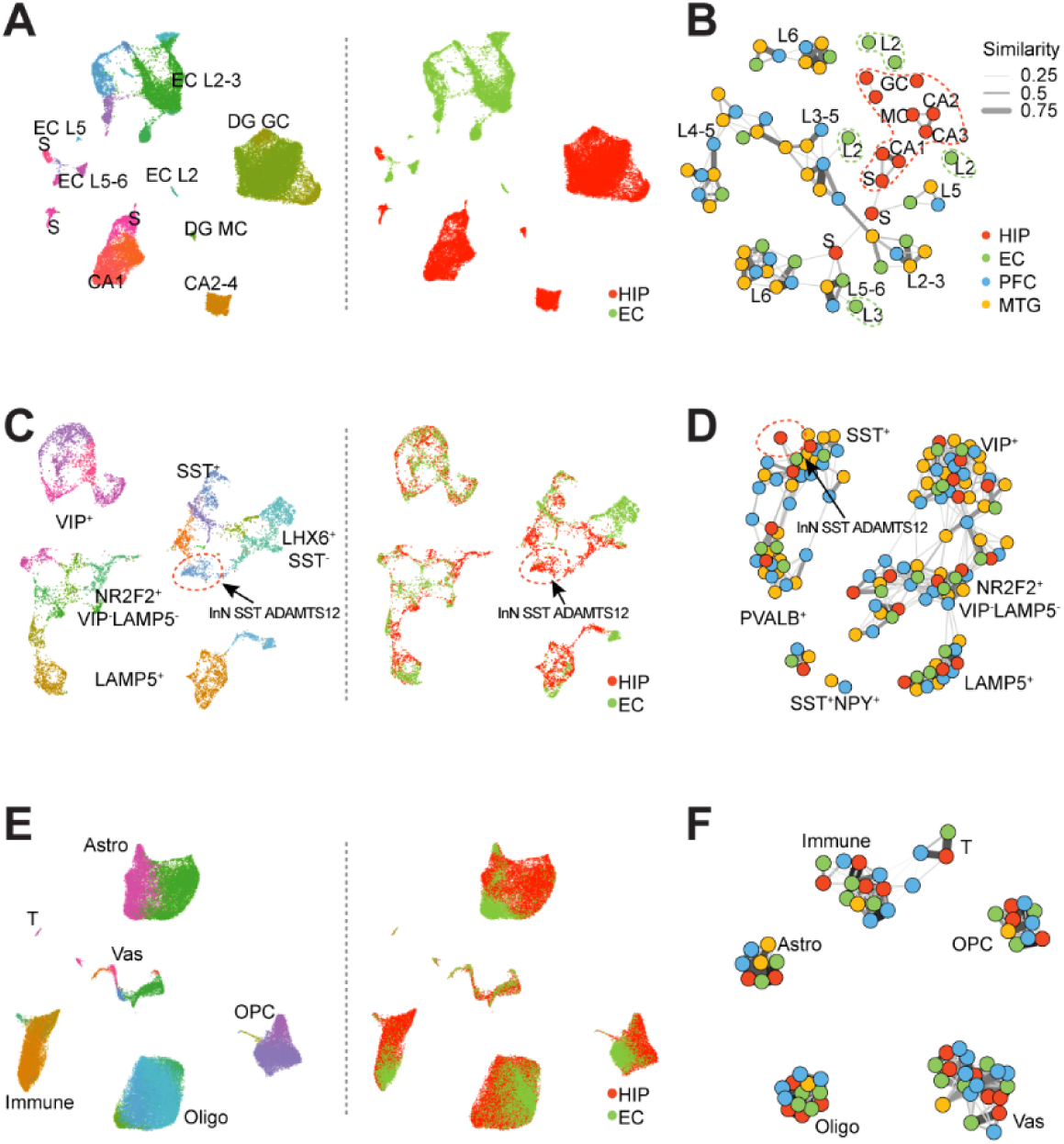
Transcriptomic distinction and similarity of hippocampal and entorhinal cell types. **A,** Left: UMAP embedding showing all excitatory neuronal subtypes detected in the hippocampal-entorhinal system, with naming conventions as in Figure 1. Right, as in left panel, but spatially colored according to two major segregated regions. HIP, hippocampus formation including dentate gyrus (DG), CA1-4 and subiculum (S); EC, entorhinal cortex. **B,** Network demonstrating the extent of transcriptome similarities among excitatory neuronal subtypes of HIP, EC, medial temporal gyrus (MTG) and dorso-lateral prefrontal cortex (dlPFC). Dots represent the subtypes within each brain region and the widths of lines represent the strength of similarity. Region, subregion and subtype information is indicated by colors and notes. Distinct subtypes of HIP and EC were outlined in corresponding colors. **C-F**, As in panels **A-B,** for inhibitory neurons (**C, D**) and non-neuronal cells (**E, F**). See also Figure S2.

**Figure 3.**
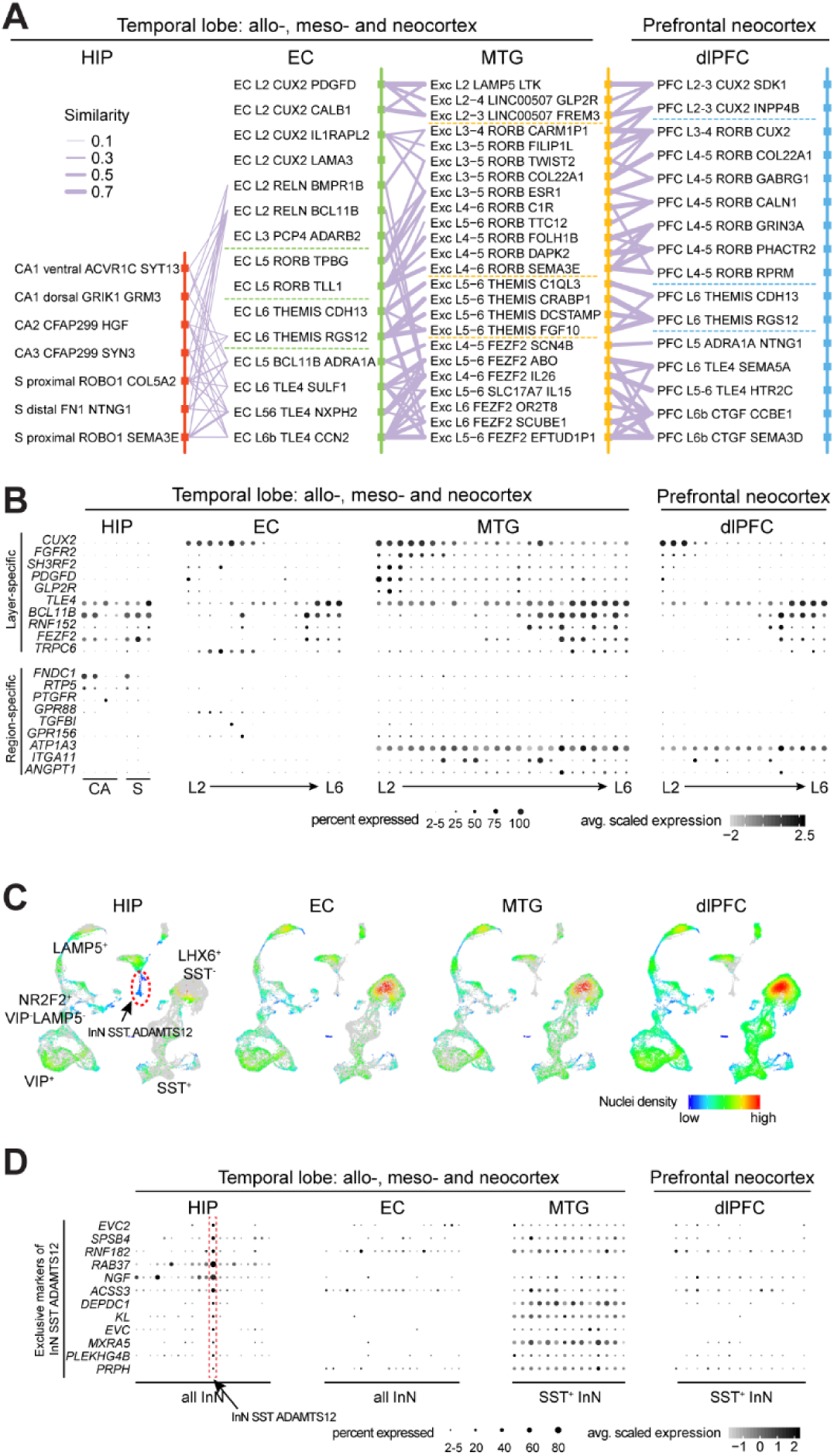
Taxonomic relationships across allo-, meso- and neo-cortex. **A,** Transcriptomic relations across subtypes of pairwise regions organized according to layer distributions. The widths of lines denote the strength of the similarities. Broad layer distinction was marked by dotted lines. **B,** Expression of neocortical upper-layer (*PDGFD*, *GLP2R*, *SH3RF2*, *FGFR2* and *CUX2*) and deep-layer markers (*TLE4*, *FEZF2*, *BCL11B*, *RNF152* and *TRPC6*) across subtypes of each region, as well as region-specific genes across these regions. The size and color of dots indicate the percent of expressed nuclei and the average gene expression within each subtype, respectively. **C,** UMAP layout exhibiting the relative density distribution of nuclei across regions. Nuclei from each region were labeled in rainbow colors to indicate density while nuclei from other regions were colored in grey. Hippocampus-enriched interneuron cluster ‘InN SST ADAMTS12’ is outlined in the plots. **D,** Expression of the exclusive markers in cluster InN SST ADAMTS12 across all inhibitory neuron subtypes in HIP and EC and all SST^+^ inhibitory neuron subtypes in MTG and dlPFC. See also Figure S3.

In particular, we identified three major transcriptomically-defined subtypes of excitatory neurons within subiculum (S), two within CA1, and only one predicted within CA2 and CA3, which is consistent with previous single cell RNA-seq studies in rodents (Zeisel et al., 2015; Cembrowski et al., 2016b) and evidence of the laminar organization of pyramidal (excitatory neurons) within the CA fields (Nielsen et al., 2010; Slomianka et al., 2011; Cipriani et al., 2016). Moreover, we found that deep-layer excitatory neuron subtypes in the neocortex were well-represented in the EC and to a lesser extent in the HIP, but upper-layer neuron subtypes were not well represented (Fig 3A-B **and** S3A). For example, we identified two *RELN*-expressing layer 2 subclusters in the EC that, similar to a previous report (Witter et al., 2017), did not correspond closely to any excitatory neuron subtype detected in the neocortex. Consistent with this observation, molecular markers for deep-layer excitatory neurons in the neocortex displayed higher expression in each subtype of HIP as compared to upper-layer molecular markers (Fig. S3B). Moreover, we observed lower expression of key molecular markers of intratelencephalic (intracerebral) projection neurons in each of the HIP excitatory neuron subtypes as compared to other neocortical neuron populations. Fig. S3C), which may be relevant to the previous observation that HIP in rodents CA fields don’t have callosally projecting intratelencephalic/intraceberal excitrory neurons (Cenquizca and Swanson, 2007). Several key genes expressed by excitatory neurons, including *FNDC1*, *RTP5*, and *PTGFR*, also exhibited specificity for allo-, meso-, or neo-cortex.

In contrast, neither inhibitory interneurons nor non-neuronal cells exhibited an obvious transition between allo-, meso-, and neo-cortex similar to that observed for excitatory neurons, with just one *SST*-expressing interneuron population in the hippocampus (InN *SST ADAMTS12*) lacking a clear counterpart in EC, MTG, and dlPFC (Fig. 2D **and** 3C). Cells in this hippocampal-specific interneuron population were notable for their expression of two EvC Ciliary Complex genes, *EVC* and *EVC2* (Caparros-Martin et al., 2013) (Fig. 3D), which may play a role in hippocampal ciliary sonic hedgehog signaling (Breunig et al., 2008; Rhee et al., 2016; Park et al., 2019). Lastly, non-neuronal cell types constituted the most transcriptomically conservative populations across the allo-, meso-, and neo-cortical taxonomy, with a high similarity observed in each subtype across all regions (Fig. 2E-F **and** S2E-J). Taken together, these finding indicate that most prominent differences across allo-, meso-, and neo-cortex occur among excitatory neurons, including the increased prevalence of intratelencephalic projection neurons in the neocortex as compared to allocortex.

### Transcriptomic insights into adult hippocampal neurogenic capacity

Neurogenesis of granule cells in the adult DG has been extensively studied in rodents and documented in many mammalian species. Many of these studies investigate the presence of cells expressing DCX, a marker of immature neurons, as a reliable indicator of newly generated neurons in DG (Couillard-Despres et al., 2005; Patzke et al., 2015; Kempermann et al., 2018). However, there is no consensus regarding the existence of significant neurogenesis in the adult human DG. Previous studies have provided evidence for the generation of granule cells in the adult and aged human DG through the detection of cell proliferation (Eriksson et al., 1998; Spalding et al., 2013), and a recent study reported a prominent population of DCX-expressing cells in the adult human DG (Moreno-Jimenez et al., 2019). Consistent with these observations, *DCX* gene expression is detected in the adult and aged human HIP, albeit dramatically lower than in the developing human or adult macaque HIP (Sousa et al., 2017; Zhu et al., 2018). Conversely, other studies have directly challenged this conclusion, having failed to identify neural progenitors or DCX-expressing granule cells after childhood in the adult human DG (Dennis et al., 2016; Cipriani et al., 2018; Sorrells et al., 2018). To add insight to these controversial sets of observations, we investigated our snRNA-seq data set to identify cells that may express *DCX* and other key gene markers related to proliferation and early neuronal differentiation that were previously characterized in adult DG neurogenesis.

Although we observed moderate expression of *DCX* in some excitatory neurons, and generally greater expression in many interneuron subtypes across the hippocampal-entorhinal system (Fig. 4A-B, S4A), we did not identify discrete clusters of *DCX*-expressing cells in HIP. However, we found 125 cells within the cluster of 14,703 DG granule cells with at least one *DCX* mRNA molecule (UMI ≥ 1) (Fig. 4C). To further characterize these *DCX*-expressing cells, we assessed whether they were enriched for markers indicative of intermediate progenitor cells and immature granule cells, the cell types previously shown to express *Dcx* during adult DG neurogenesis (Couillard-Despres et al., 2005; Patzke et al., 2015; Kempermann et al., 2018). Although some of these cells co-expressed migrating or immature neuron markers including *NEUROD2* (61.6%), *FXYD7* (32.8%), and *NCAM1* (97.6%) (UMI ≥ 1) (Hochgerner et al., 2018) (Fig. 4E), we found that these percentages were generally comparable or lower than those from excitatory neurons of EC (51.9%, 61.9%, and 99% for *NEUROD2*, *FXYD7*, and *NCAM1*, respectively) (Fig. 4D, F), a region where adult newborn neurons have not been reported to be generated. Moreover, we did not observe enrichment for these markers as compared to *DCX* non-expressing DG granule cells (nominal p-value > 0.05) (Fig. S4A). Similarly, putative neural progenitor cells expressing *MKI67*, which precede immature neurons in development, did not constitute an independent cluster in DG (Fig. S4B). We also found no evidence that neural stem cells clustered with astrocytes, a transcriptomically similar cell population, as all *NES*-expressing cells in the astrocytic cluster co-expressed *AQP4* (61 of 61 cells), a marker for differentiated cells committed to the astrocytic lineage (Fig. S4C). Far fewer of these cells expressed *HOPX* (8 of 61 cells), a reported marker for quiescent progenitors responsible for adult neurogenesis in mouse (Berg et al., 2019).

**Figure 4.**
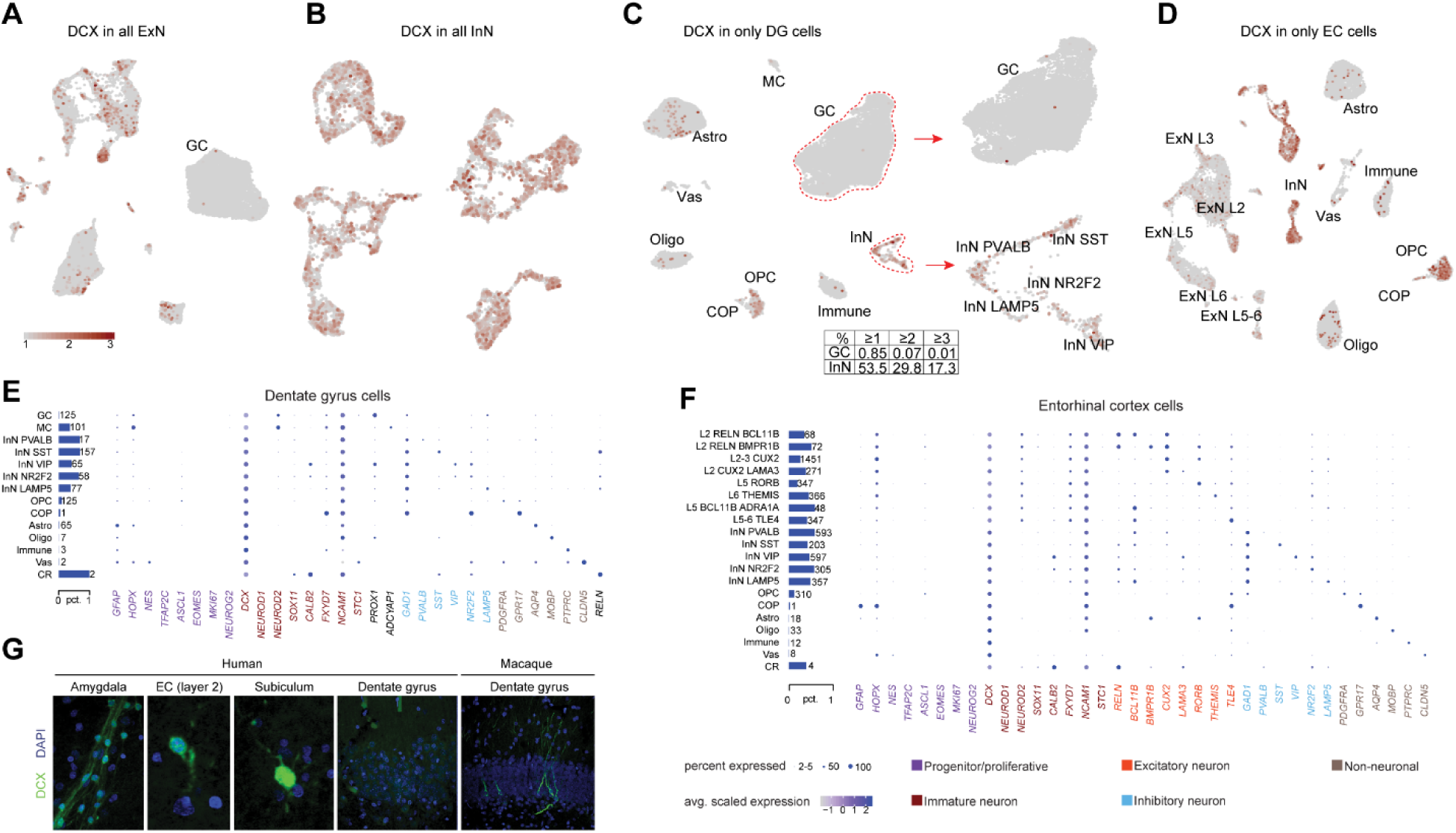
DCX expression in hippocampal and entorhinal cell types. **A,** Expression of *DCX* in all excitatory neurons visualized in UMAP embedding. Granule cells (GC) with sparse *DCX* expression are noted. **B,** Expression of *DCX* in all inhibitory neurons visualized in UMAP embedding. **C,** *DCX* expression in only dentate gyrus (DG) cells displayed in UMAP embedding. Major cell type classes are labeled. Zoom-in views illustrate the expression of *DCX* specifically in four subtypes of inhibitory neurons and GC. Bottom table denotes the percentages of GC and InN expressing *DCX* over UMI thresholds of 1, 2 and 3. **D,** *DCX* expression in only EC cells displayed in UMAP embedding. Cell types expressing *DCX* (red) are labeled. **E,** Bar plot shows the numbers and percentages of *DCX*+ cells within each cell type of DG. Dot plot shows the expression of markers of neural stem cells (NSC), neural progenitor cells (NPC), migrating and immature neurons (IM), as well as markers labeling different cell types in *DCX*+ cells of DG. The size and color of dots indicate the percent of expressed nuclei and the average gene expression within each type, respectively. **F**, Bar plot shows the numbers and percentages of *DCX*+ cells within each cell type of EC. Dot plot shows the expression of markers of neural stem cells (NSC), neural progenitor cells (NPC), migrating and immature neurons (IM), as well as markers labeling different cell types in *DCX*+ cells of EC. The size and color of dots indicate the percent of expressed nuclei and the average gene expression within each type, respectively. **G,** Images of immunofluorescent cells expressing DCX in regions of human and macaque brain. See also Figure S4 and Table S1.

We complemented these snRNA-seq analyses using immunohistochemistry with two different commonly used antibodies against DCX. As recently reported (Sorrells et al., 2019), we detected many DCX-immunopositive neurons in the paralaminar nuclei of the adult human amygdala (Fig. 4G). In contrast, although we detected *DCX* transcripts in all brains processed for snRNA-seq, immunohistochemical detection of DCX in the hippocampal-entorhinal system was successful in less than one-third of an independent cohort of postmortem adult human samples (n=11; **Table S1**). This included scarce DCX-immunopositive neurons in the subiculum that were weakly immunopositive for the inhibitory neuron marker GAD1, and the EC, which were not GAD1-immunopositive (Fig. 4G). Moreover, we were unable to detect DCX-immunopositive cells in the DG or the adjacent CA4 field, including in one brain also used for snRNA-seq (total of 12 brains), indicating a poor correlation between detection of DCX mRNA and protein in the postmortem adult human samples. Together, these findings are consistent with recent DCX immunohistochemical studies (Dennis et al., 2016; Cipriani et al., 2018; Sorrells et al., 2018) showing that neurogenesis does not continue, or is extremely rare, in the adult human DG.

### Species, age and excitatory neuron subtype-specific *METTL7B* expression

Tissue and single cell expression profiles, including from multiple subregions of the HIP and EC, allowed us to integrate regional and cell type-specific differences in disease susceptibility with temporal patterns of gene expression. We began by identifying candidate genes enriched in the hippocampus and whose expression changes with age as described in a developmental and multi-regional human brain transcriptome dataset we previously generated (Kang et al., 2011; Li et al., 2018). Genes were ranked based on their region-specific changes in expression over development and aging (Fig. 5A **and Table S2**). Within the hippocampus, of the three genes exhibiting the greatest increased (*KL, METTL7B, PTGS1*) or decreased expression (*TSHR, MSTN, WNT8B*) across time, five have been previously functionally characterized to some extent, and the two that exhibited a progressive increase (*KL* and *PTGS1*) have been associated with Alzheimer’s disease (Qin et al., 2003; Zeldich et al., 2014).. In contrast, the second-ranked upregulated gene, *METTL7B*, has not been comprehensively studied in the context of the vertebrate brain. *METTL7B*, which is predominantly expressed in liver (Uhlen et al., 2015), encodes a membrane protein associated with endoplasmic reticulum (ER) and lipid droplets, and, by amino acid sequence homology, is predicted to belong to the protein methyltransferase superfamily (Turro et al., 2006; Thomas et al., 2013). We confirmed that at the RNA and protein level, METTL7B is enriched in the adult human hippocampus (Fig. 5B-D). Analysis of *METTL7B* in the same 16 homologous brain regions in chimpanzee and rhesus macaque (Sousa et al., 2017; Zhu et al., 2018) found that expression in the hippocampus, and the cerebrum more generally, is not distinct to humans. However, *METTL7B* was more broadly expressed throughout the cerebrum in rhesus macaque brain (Fig. S5A). The expression of *METTL7B* was also enriched in the human and macaque cerebrum as compared to the cerebrum of mouse, rat, rabbit, and opossum (Cardoso-Moreira et al., 2019) (Fig. S5B).

**Figure 5.**
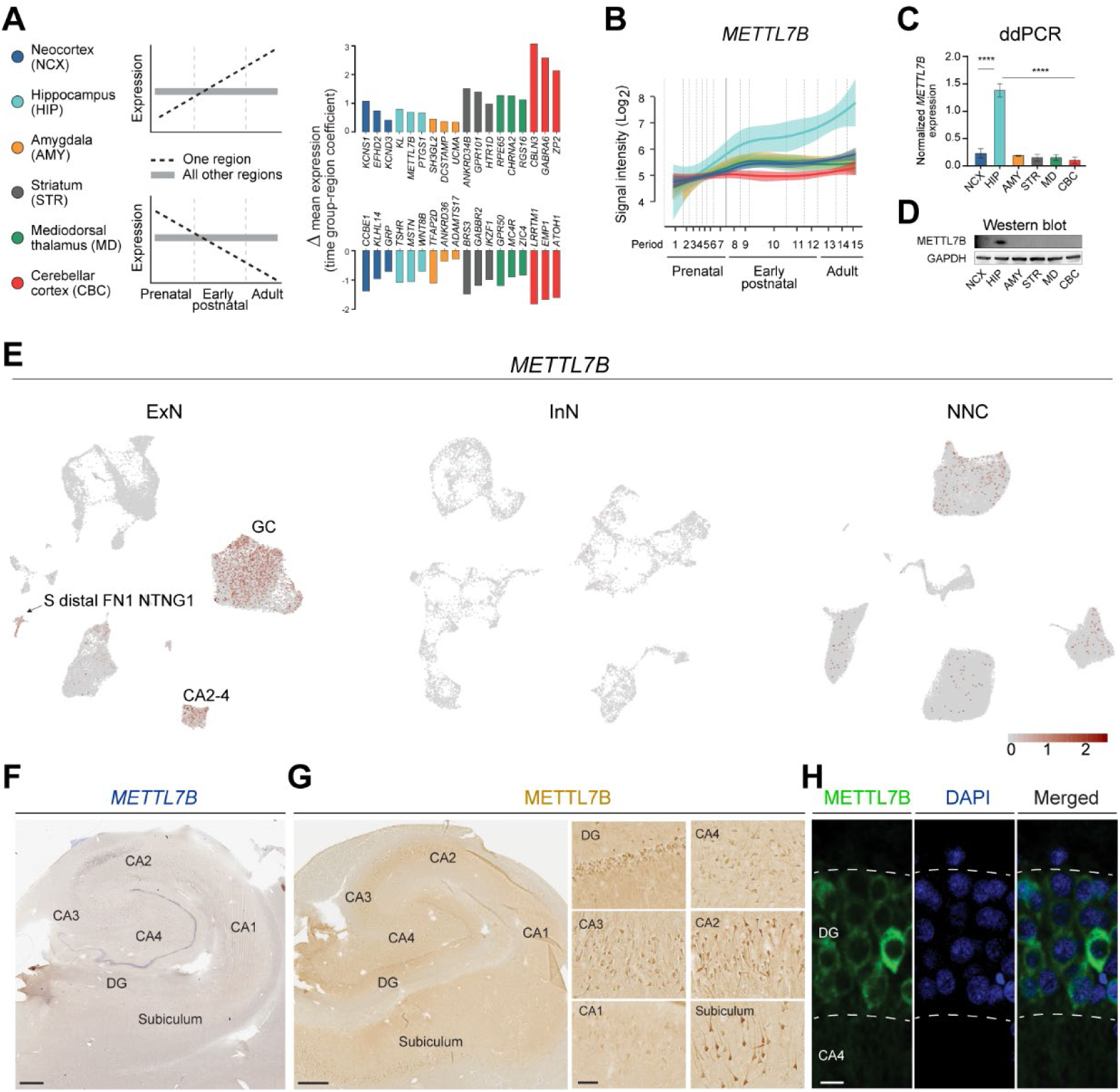
Human METTL7B is enriched in dentate gyrus granule cells. **A**, Top three up- and down-regulated genes in six brain regions throughout development. **B**, Expression trajectory of METTL7B showing enrichment in hippocampus and an increasing expression with age. **C-D**, Digital droplet PCR and immunoblot validation in six regions of adult brain. One-way ANOVA with post-hoc Dunnett’s adjustment (****P<0.0001), N=3 per group. **E**, *METTL7B* expression in GCs displayed in UMAP embedding. **F-H**, *In situ* hybridization and immunostaining of adult hippocampus show prominent labeling of dentate gyrus granule cells, and subicular and CA2 pyramidal neurons. Scale bars = 1 mm; insets = 100 μm; immunofluorescence = 10 μm. See also Figure S5 and Table S2.

We next mapped the cell type expression of *METTL7B*, and found it is virtually exclusive to excitatory neurons, with highest enrichment in the DG, followed by CA2-3 and then subiculum (Fig. 5E). RNA *in situ* hybridization and immunolabeling of adjacent sections confirmed that the highest signal intensity was in DG granule neurons and pyramidal neurons in CA2, with lesser expression in CA3-4 subfields and Sub. (Fig. 5F-H). Prompted by the cortical cell-type taxonomic similarities we described above, we also analyzed *METTL7B* expression in the neocortex and found high levels in the large pyramidal neurons of layer 5B (Fig. S5D), such as Betz and Meynert cells in M1C and V1C, respectively. Interestingly, these subregions/layers and cells are generally among the last to exhibit hallmarks of Alzheimer’s disease pathology including the formation of plaques, tangles, and neuronal death (McMenemey, 1940; Wilcock and Esiri, 1982; Hof and Morrison, 1990; Davies et al., 1992; Jin et al., 2004; Schonheit et al., 2004; West et al., 2004; Ohm, 2007; Braak and Del Trecidi, 2015). Immunostaining of adult hippocampal and neocortical tissue sections of rhesus macaque, a widely studied Old World monkey, revealed staining profiles comparable to those of humans, with hippocampal DG, CA2 and subicular pyramidal neurons, as well as neocortical large L5B pyramidal neurons, displaying strong immunolabeling for METTL7B (Fig. S5E-F). By contrast, we observed very little expression of METTL7B in cortico-cortical pyramidal neurons of neocortical and entorhinal L5A and upper layers (L2-4), as well as hippocampal CA1 pyramidal neurons (Fig. 5G, Fig. S5C-E), which are known to be selectively vulnerable in aging and the initial stages of Alzheimer’s disease in humans (McMenemey, 1940; Wilcock and Esiri, 1982; Hof and Morrison, 1990; Davies et al., 1992; Jin et al., 2004; Schonheit et al., 2004; West et al., 2004; Ohm, 2007; Braak and Del Trecidi, 2015). We also found that the homolog of *METTL7B* is not expressed in adult mouse brain (Fig. S5F-I), indicating that METTLB’s expression and function in the adult brain is species-specific.

### METTL7B interacts with Alzheimer’s disease-related proteins

To identify METTL7B interacting proteins, we performed unbiased proteomic analysis using two different affinity-based approaches to find METTL7B interacting partners. The first approach utilized HaloTag fusion protein technology (Fig. S6A) and has scarce non-specific binding (Hook, 2014). We created stable cell lines by transducing a human cortical neural progenitor cell line, which has been previously utilized to model Alzheimer’s disease-related molecular processes (Choi et al., 2014), to express either HaloTag or METTL7B-HaloTag fusion protein. Captured proteins, representing proteins putatively interacting with METTL7B-HaloTag, were detected by silver stain (Fig. S6B) and analyzed by LC-MS/MS (**Table S3**). We used co-immunofluorescence to observe a high degree of overlap of METTL7B with CALNEXIN and ADFP, markers of the ER and lipid droplets (LD), respectively (Turro et al., 2006) (Fig. S6C). Using Significance Analysis of INTeractome (SAINT) (Choi et al., 2011), we identified 275 true METTL7B interactors (Fig. S6D**, Table S4**). Fold-enrichment analysis for major subcellular compartments revealed these true METTL7B interactors showed significant enrichment in ER- and LD-associated proteins (Fig. S6E). KEGG pathway enrichment analysis revealed the highest enrichment in the proteasome, protein processing in the ER, oxidative phosphorylation, and three neurodegenerative diseases: Parkinson’s disease, AD, and Huntington’s disease (Fig. S6F).

We next pursued an overlapping approach using BioID technology, which utilizes biotin ligase (BirA) activity to biotinylate proteins in the vicinity of a protein of interest and better capture interactions that may be weak or transient (Fig. S6G-H**, Table S3**) (Roux et al., 2012). Here again, METTL7B interactors co-localized with molecular markers for the ER and lipid droplets (Fig. S6I). We identified 1794 true METTL7B interactors that were enriched in ER and lipid droplets-associated proteins (Fig. S6J-K**, Table S4**). Molecular pathways involving true interactors were related to endocytosis, SNARE interactions in vesicular transport, and protein-protein processing in ER (Fig. S6L). Moreover, several putative interacting proteins associated with METTL7B have been previously implicated in Alzheimer’s disease in a variety of ways, such as: amyloid precursor protein (APP), inhibition of γ-secretase (RTN3 and RTN4/NOGO), amyloid binding (NAE1, LRP1, APBB1), protein (de)phosphorylation (PPP3CA, UQCRFS1, CDK5, CALM1, EIF2AK3), APP proteolytic cleavage (ADAM17, CASP3), mitochondrial function (COX6B1, CYCS, ATP5B), calcium homeostasis (ITPR1, ITPR2), ER stress monitoring (ATF6), and protein folding (PDIA3). We confirmed that many of these genes were extensively co-expressed with *METTL7B* in several hippocampal populations (Fig. S7).

Intersecting the lists of putative METTL7B interacting proteins from both strategies, we found 100 high-confidence proteins, with the most enriched gene ontology term being protein processing in ER (Fig. 6A-B). To further explore the functional significance of the METTL7B interaction network, we examined the profile of spatial overlap between METTL7B true interactors and all KEGG pathways in protein-protein interaction networks, and evaluated the significance of overlap as a function of network distance (Fig. 6C-D). We found that the highest-confidence METTL7B true interactors overlap with several neurodegenerative diseases curated in KEGG pathways in the most proximal network distance from METTL7B, including KEGG Alzheimer’s Disease Pathway and Protein Processing in ER (Fig. 6E). We subsequently used immunoblots to confirm that candidate proteins RTN4, APP, and LRP1 were specific to METTL7B sample eluates. RTN3 was not detected by immunoblot in any of the samples, possibly due to low pull-down amounts that were below Western blot detectability threshold (Fig. 6F-G). We next sought to determine whether these proteins were potential substrates for METTL7B methyltransferase activity. To do so, we incubated purified recombinant proteins in a continuous enzyme-coupled S-adenosylmethionine (SAM) methyltransferase assay, which responds fluorescently to S-adenosylhomocysteine, a byproduct of SAM methyltransferase activity. All four assayed samples in which candidate proteins were incubated with recombinant METTL7B produced a significant increase in signal compared to candidate proteins incubated alone (Fig. 6H). These results suggest that METTL7B uses SAM as a methyl donor, and that METTL7B may act enzymatically on proteins specifically related to Alzheimer’s disease pathology.

**Figure 6.**
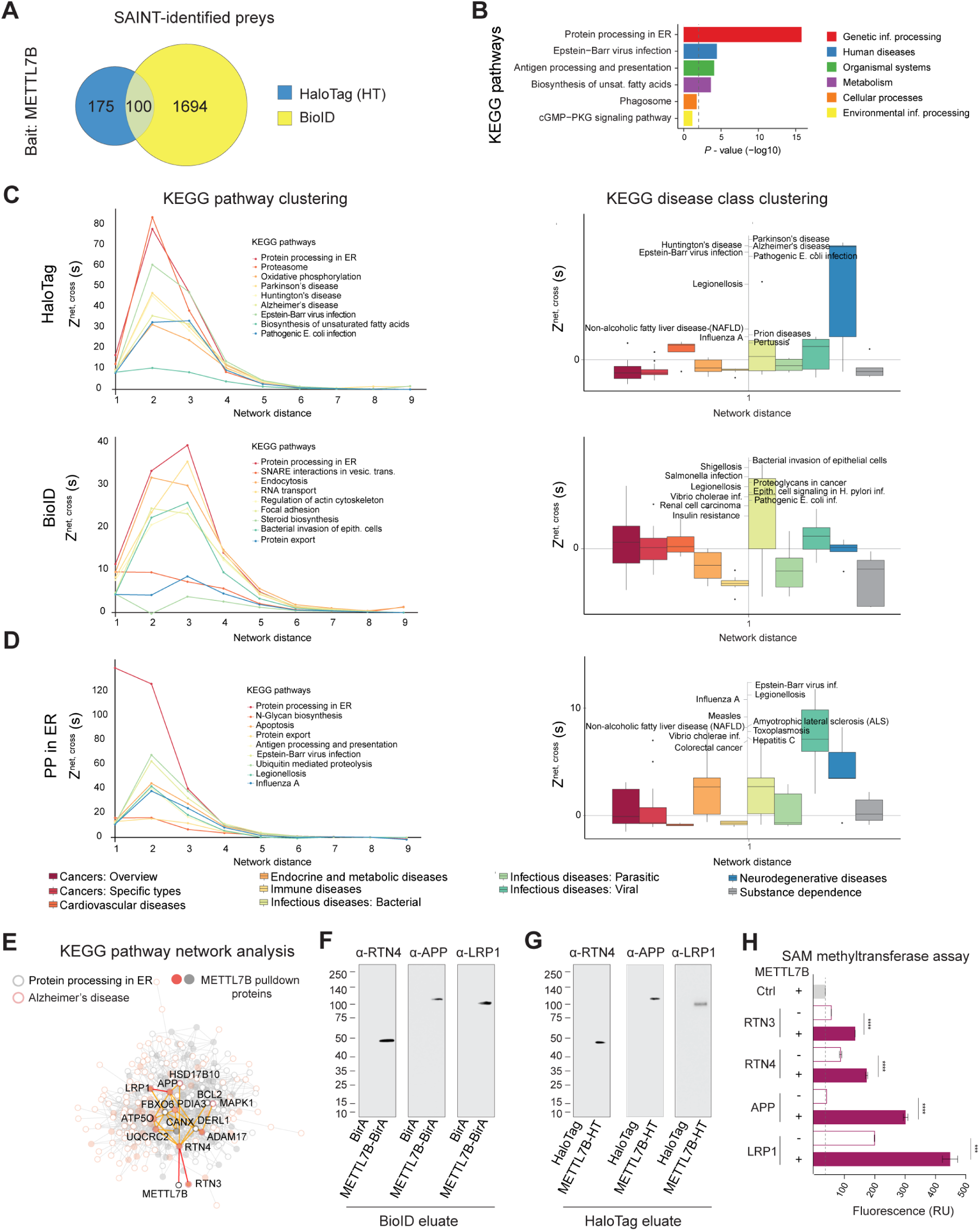
METTL7B-interacting proteins are enriched in the ER and are associated with AD. **A**, Venn diagram of true METTL7B interacting proteins. **B**, Protein processing in ER (PPER) is the most significantly enriched KEGG pathway of METTL7B interactors. **C**, Comparing BioID to HaloTag, the significance of Z^net, cross^ peak shifts toward a closer network distance (left panel). At the closest network distance, the biggest shift in the significance of Z^net, cross^ occurs for several neurodegenerative diseases: Parkinson’s disease, AD, and Huntington’s disease (right panel). **D**, Validation of spatial clustering analysis using PPER against all KEGG pathways. As expected, PPER is the strongest overlapping KEGG pathway. **E**, Subset of protein-protein interaction network with all the proteins in the KEGG PPER (grey empty circle) as well as those in Alzheimer’s disease pathway (orange empty circle). True METTL7B interactors are shown as filled circles. Top candidate proteins as well as their direct protein-protein interacting partners are highlighted. **F-G**, Immunoblot confirmation of top candidates. **H**, SAM-assay showing an increase in methylation in the presence of METTL7B. *P*-values calculated by unpaired two-tailed Student’s t test, N=3. All data are mean ± SEM. \*\*\*\**P*< 0.0001, \*\*\**P*< 0.001. See also Figures S6, S7, and Tables S3 and S4.

### METTL7B reduces amyloid beta generation

Given the protein interactions and potential activity of METTL7B on APP, we next sought to determine whether METTL7B has an effect on APP proteolytic processing, which plays an important role in Alzheimer’s disease pathology (Hardy and Selkoe, 2002; Tanzi and Bertram, 2005; Ballatore et al., 2007). We therefore transduced N2a cells, a mouse neuroblastoma cell line that does not natively express endogenous *Mettl7b* (Fig. 7A), to express full length human APP_695_ along with METTL7B or EGFP. As determined by an enzyme-linked immunosorbent assay (ELISA), METTL7B expression resulted in a ∼31.8% mean reduction in Aβ_40_ and a ∼34.9% mean reduction in Aβ_42_ levels, with no significant change in Aβ_42_/Aβ_40_ ratio (Fig. 7B-C). However, we found that the availability of METTL7B for putative interactions with Alzheimer’s disease-associated proteins may be limited in conditions with high levels of lipids, as supplementing cell culture media with linoleic and oleic acid complexes resulted in the increased formation of lipid droplets along with a commensurate shift of METTL7B from the ER to lipid droplets, but not METTL7B-interacting and Alzheimer’s disease-associated proteins (Fig. 7D-E).

**Figure 7.**
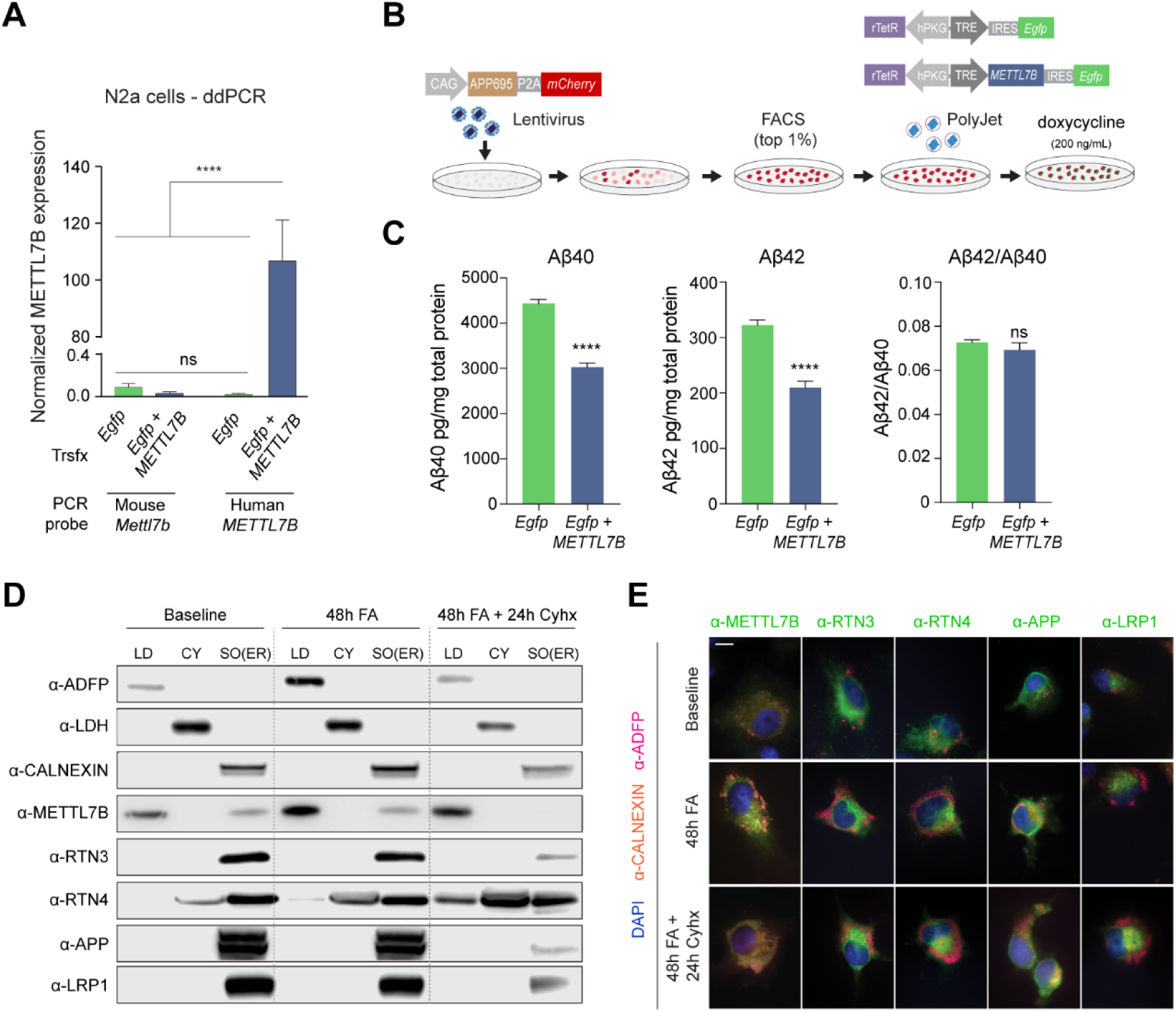
METTL7B reduces Aβ generation. **A**, ddPCR of human *METTL7B* misexpression in Neuro-2a cells. One-way ANOVA with post-hoc Dunnett’s adjustment, N=3 per group. **B**, Schematic of experimental design. **C**, ELISA for Aβ_40_ and Aβ_42_ levels in cells with or without *METTL7B*. *P*-values calculated by unpaired two-tailed Student’s t-test, N=6 per group. All data are mean ± SEM. \*\*\*\**P*<0.0001. **D-E**, Immunoanalysis of human neural cells. Increased fatty acid load leads to shift of METTL7B from ER to LDs, while Alzheimer’s disease-associated high confidence interactors remain unaffected. Blocking translation of new proteins with cycloheximide (Cyhx) suggests a complete shift of METTL7B. Scale bar = 10 μm. CY = cytosol; SO = sedimentable organelle.

## DISCUSSION

Here we present a detailed single cell transcriptomic analysis of cells in the adult human hippocampal-entorhinal system, and through analysis of this resource describe novel biology related to the molecular diversity of these cells. We found region-specific distinctions in gene expression patterns, particularly of excitatory neurons, with clear implications for hippocampal-entorhinal physiology.

These data and accompanying analyses refine our understanding of the evolution of allo-, meso-, and neo-cortex. The transcriptomic signatures we developed strongly suggest homology between mammalian allocortex and specifically deep layers of the EC and neocortex. Analyses of the transcriptome also suggest that adult neurogenesis in the DG is, if present, extremely limited, with no clear signatures of newborn or immature excitatory neurons present. Although this is not necessarily in conflict with previous efforts, some of which have detected newborn neurons in some but not all adult human hippocampal specimen (Spalding et al., 2013), our analysis of both DCX RNA and protein suggest further analyses is necessary to discern the neurogenic capacity of the adult human hippocampus.

In addition, we implicated METTL7B in aging and disease processes of the human hippocampus. We found strong evidence that METTL7B is enriched in subregions/layers and cell types known to be less vulnerable to initial Alzheimer’s disease neuropathology in the HIP, EC and primary neocortical areas. We also found that METTL7B interacts with important Alzheimer’s disease-related proteins (e.g., APP, LRP1, RTN3, and RTN4), and that its overexpression reduced Aβ levels. In addition, our results also suggest that the subregion/cell type enrichment we observed could be Catarrhini (i.e., the Old World monkeys and the apes)-specific, supporting the contention that manifestation of some Aβ plaque pathology may have distinct manifestation in primates (Rapoport, 1989). Moreover, brain regions and circuits preferentially affected in early stages of Alzheimer’s disease are generally expanded in humans, other apes, and Old World monkeys (Stephan, 1975; Demeter et al., 1990; Morrison and Hof, 1997; Suzuki and Amaral, 2004; Braak and Del Trecidi, 2015), and aged macaques and great apes exhibit early signs of Alzheimer’s disease-like amyloid and Tau pathology (Perez et al., 2013; Finch and Austad, 2015; Edler et al., 2017; Paspalas et al., 2018).

Our investigation of METTL7B may also lend support to evidence suggesting the role of several classes of lipids and lipid droplets in Alzheimer’s disease pathogenesis. This evidence stems from knowledge of the APOE4 allele, encoding a cholesterol carrier, that is associated with an increased risk for late onset Alzheimer’s disease (Strittmatter et al., 1993). In addition, cholesterol dyshomeostasis can directly influence the activity of Aβ-related proteases and increase Aβ production (Di Paolo and Kim, 2011), and strong epidemiological evidence supports mid-life hypercholesterolemia as a risk factor for Alzheimer’s disease. Moreover, lipid droplet accumulations have been identified in postmortem Alzheimer’s disease brains, from the initial report by Alois Alzheimer to more recent reports including a transgenic mouse model of Alzheimer’s disease (3xTg-AD) (Alzheimer et al., 1995; Hamilton et al., 2015). Because METTL7B translocates to lipid droplets after fatty acid loading, its availability in the ER and consequently its ability to interact with target proteins implicated in Alzheimer’s disease pathogenesis may be diminished. Taken together, the distinct expression pattern of METTL7B in Catarrhini and the integration of METTL7B into mechanisms underlying specific cellular, regional, age, and species-specific aspects of development and disease suggests protein methylation may have previously unappreciated roles in the neurobiology of aging and disease.

## Acknowledgments

We thank K. Meyer for help with gene expression analysis; A. Rosa Campos and K. Motamedchaboki (SBP Medical Discovery Institute, La Jolla, CA) for help with generating proteomics data; M. Horn, A. Huttner, M. Pletikos, and S. Wilson for assistance with tissue acquisition and processing; and J. DeFelipe and the NIH NeuroBioBank, for providing human tissue. We also thank A. Duque for using equipment from MacBrainResource (NIH/NIMH MH113257). This work was supported by the NIH grants DA023999 (P.R.) and MH103339, MH106934, MH110926 and MH109904 (N.S.). The project that gave rise to these results received the support of a fellowship from “la Caixa” Foundation (ID 100010434). The fellowship code is LCF/BQ/PI19/11690010. Additional support was provided by the Kavli Foundation, and the James S. McDonnell Foundation.

## Author contributions

D.F. and N.S. conceived and designed the study. D.F. designed and performed most of the METTL7B experiments. J.C. analyzed proteomic data. M.S. prepared nuclei for sequencing. C.X., Q.L., S.M., and G.S. analyzed single nuclei sequencing data. A.T.N.T. performed SAM assay, digital droplet PCRs, and western blots. J.I.A., I.G., L.J.R., and A.M.M.S. contributed to additional data collection. D.F. and N.S. wrote the manuscript. All authors edited the manuscript.

## Data and materials availability

All scripts are available to investigators upon request. Supplement contains transcriptome analysis and proteomic data with analysis. Correspondence should be directed to

---

**Figure S1 (related to Figure 1).**
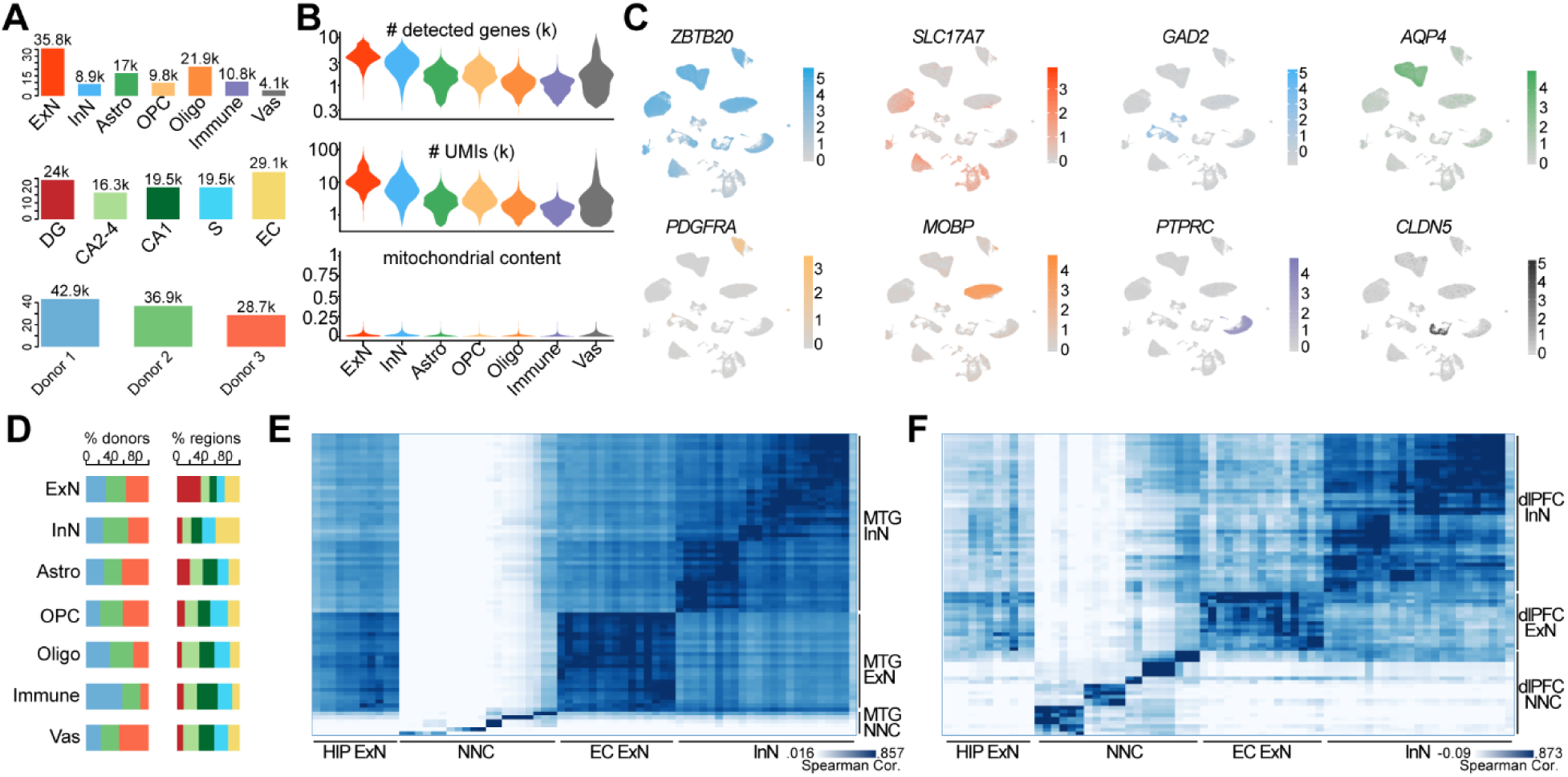
Overview of snRNA-seq data quality. **A,** Bar plots depicting numbers of nuclei passing quality control across major cell types (upper), subregions (middle) and donors (bottom). **B,** Violin plots denoting the numbers of detected genes (upper), total UMIs (middle) and mitochondrial contents (lower) across major cell types. Note the first two plots were displayed in the logarithmic scale. **C,** Expression distribution of markers of pan-hippocampus (*ZBTB20*), excitatory neuron (*SLC17A7*), inhibitory neuron (*GAD2*), astrocyte (*AQP4*), OPC (*PDGFRA*), oligodendrocyte (*MOBP*), immune cell (*PTPRC*) and endothelial cell comprising most vasculatures (*CLDN5*) projected onto the UMAP embedding. **D,** Bar plots showing donor and subregional compositions in each major cell type, with coloring scheme conforming to panel **A**. **E-F,** Transcriptome alignments between subtypes of hippocampal-entorhinal system (columns) and human medial temporal gyrus (MTG) (**E**), and human prefrontal cortex (PFC) (**F**) (rows). Cell grids in heat maps denote the Spearman correlation coefficients between each pair of subtypes across data sets.

**Figure S2 (related to Figures 1 and 2).**
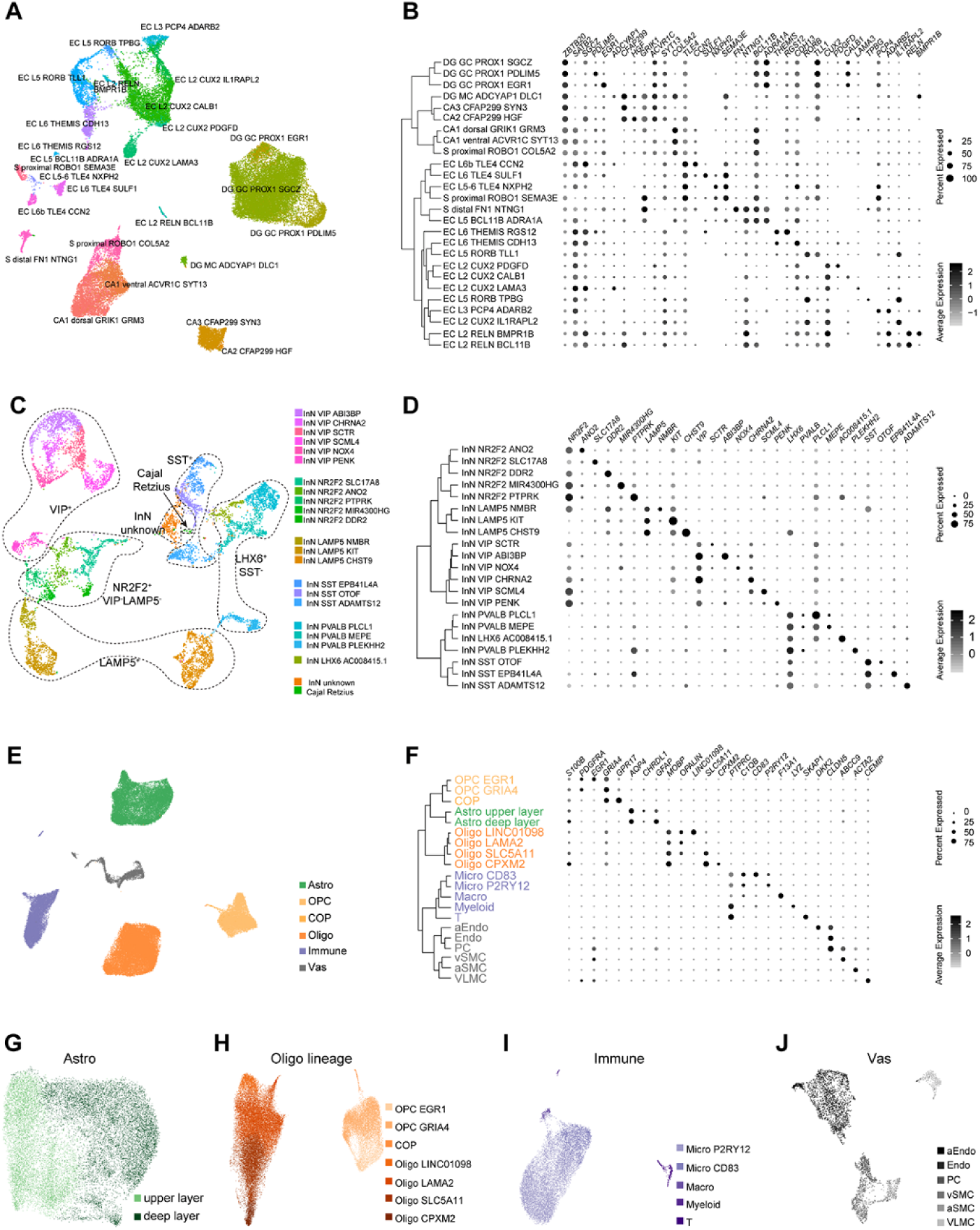
Transcriptome diversity of hippocampal and entorhinal excitatory neurons. **A,** UMAP layout showing all excitatory neuronal subtypes detected in the hippocampal-entorhinal system. **B,** Hierarchical tree describing the excitatory neuron subtypes and corresponding marker gene expression in them. The size and color of the dot plot indicate the percent of expressed nuclei and the average expression within each subtype, respectively. **C,** UMAP layout showing all inhibitory neuronal subtypes detected in the hippocampal-entorhinal system. **D,** Hierarchical tree describing the inhibitory neuron subtypes and corresponding marker gene expression in them. The size and color of the dot plot indicate the percent of expressed nuclei and the average expression within each subtype, respectively. **E,** UMAP showing all non-neuronal types detected in the hippocampal-entorhinal system. **F,** Hierarchical tree describing the non-neuronal subtypes and corresponding marker gene expression in them. The size and color of the dot plot indicate the percent of expressed nuclei and the average expression within each subtype, respectively. **G-J,** UMAP showing subtypes of astrocytes (**G**), oligodendrocyte lineage (OPCs, COPs and oligodendrocytes) (**H**), immune cells (**I**) and vasculature cells (**J**).

**Figure S3 (related to Figure 3).**
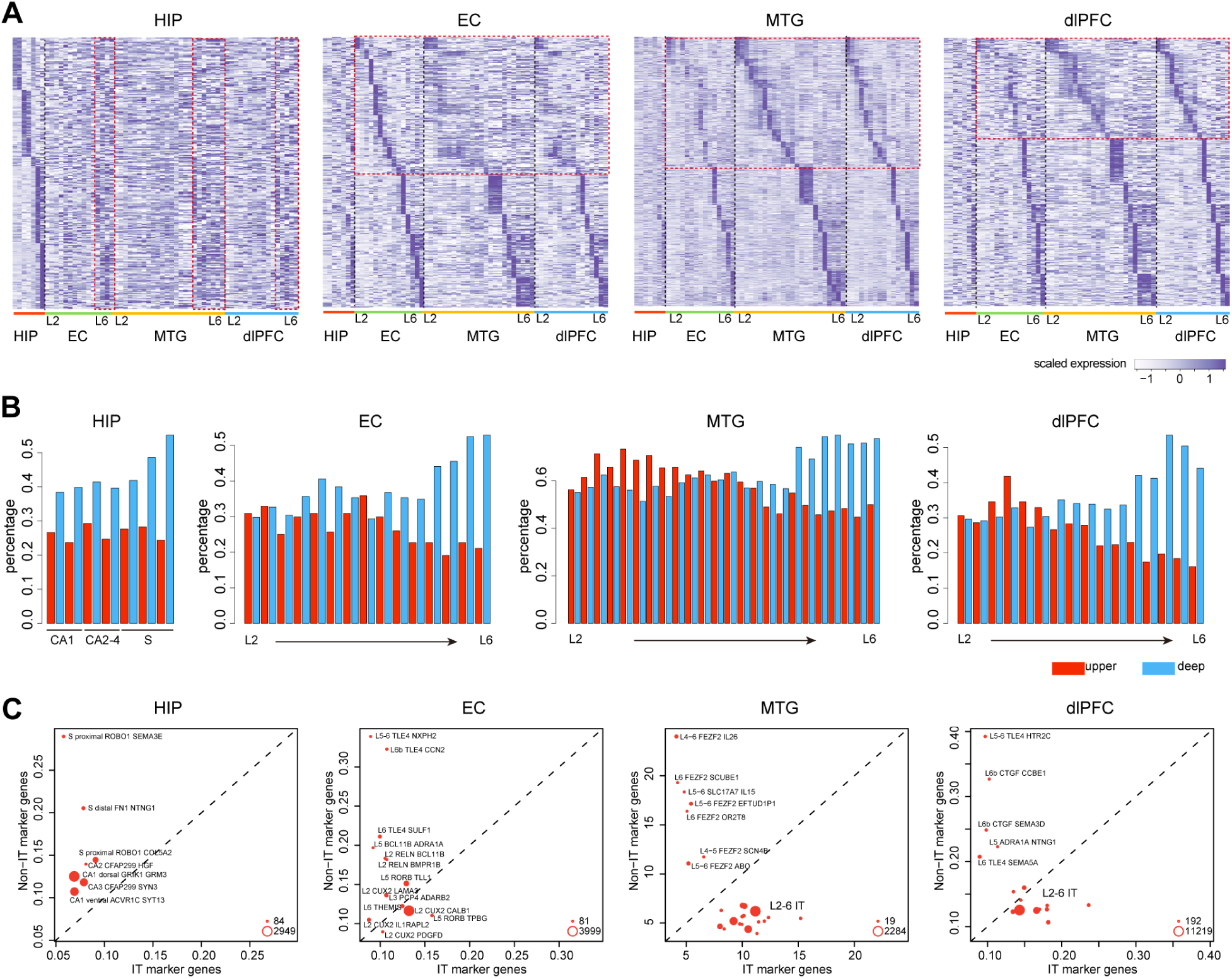
Transcriptome comparison of excitatory neurons between hippocampus, EC and neocortex. **A,** Heat map depicting the expression of marker genes from a certain region (rows) across subtypes of all the four regions: HIP, EC, MTG and PFC (columns). The relative expression enrichment of hippocampal marker genes in deep layers of EC, MTG and PFC, and the upper layer divergence between EC and neocortex are outlined. **B,** Bar plots denoting the percentages of neocortical upper-(red) and deep-(blue) layer marker genes that were expressed in each subtype of HIP, EC, MTG, and PFC. **C,** Scatter plots showing the median expression of intratelencephalic (IT) neuron markers (x axis) versus that of non-IT markers (y axis) in each subtype of HIP, EC, MTG and PFC. Dot size indicates the number of cells within the corresponding subtype.

**Figure S4 (related to Figure 4).**
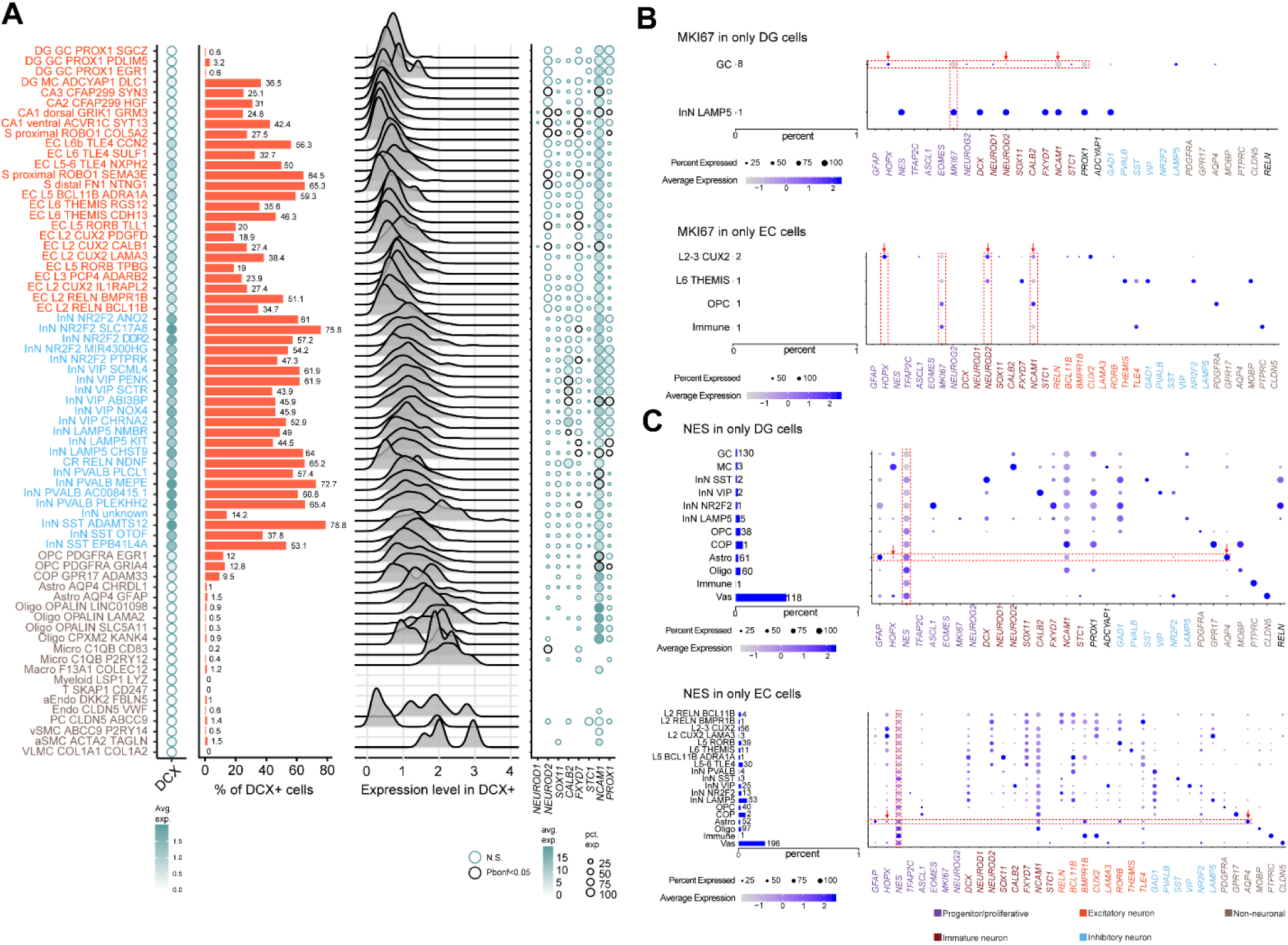
Characterization of *DCX*+, *MKI67*+ and *NES*+ cells. **A**, First panel: Average expression of *DCX* across clusters. Second panel: percentage of *DCX* positive cells in each cluster. Third panel: expression (Seurat-log normalized counts) level of *DCX*+ cells. Fourth panel: Average expression (color) and percentage of cells expressing (size) gene markers of migrating and immature granule cells, and *PROX1*. Black circle indicates significant enrichment of *DCX* and each other gene colocalization compared to *DCX*-cells calculated by means of a Fisher’s exact test. **B, C**, Bar plot shows the numbers and percentages of *MKI67*+ cells (**B**) or *NES*+ cells (**C**) within each cell type of DG (upper) and EC (bottom). Dot plot shows the expression of markers of neural stem cells (NSC), neural progenitor cells (NPC), migrating and immature neurons (IM), as well as markers labeling different cell types in *MKI67*+ or *NES*+ cells. The size and color of dots indicate the percent of expressed nuclei and the average gene expression within each type, respectively.

**Figure S5 (related to Figure 5).**
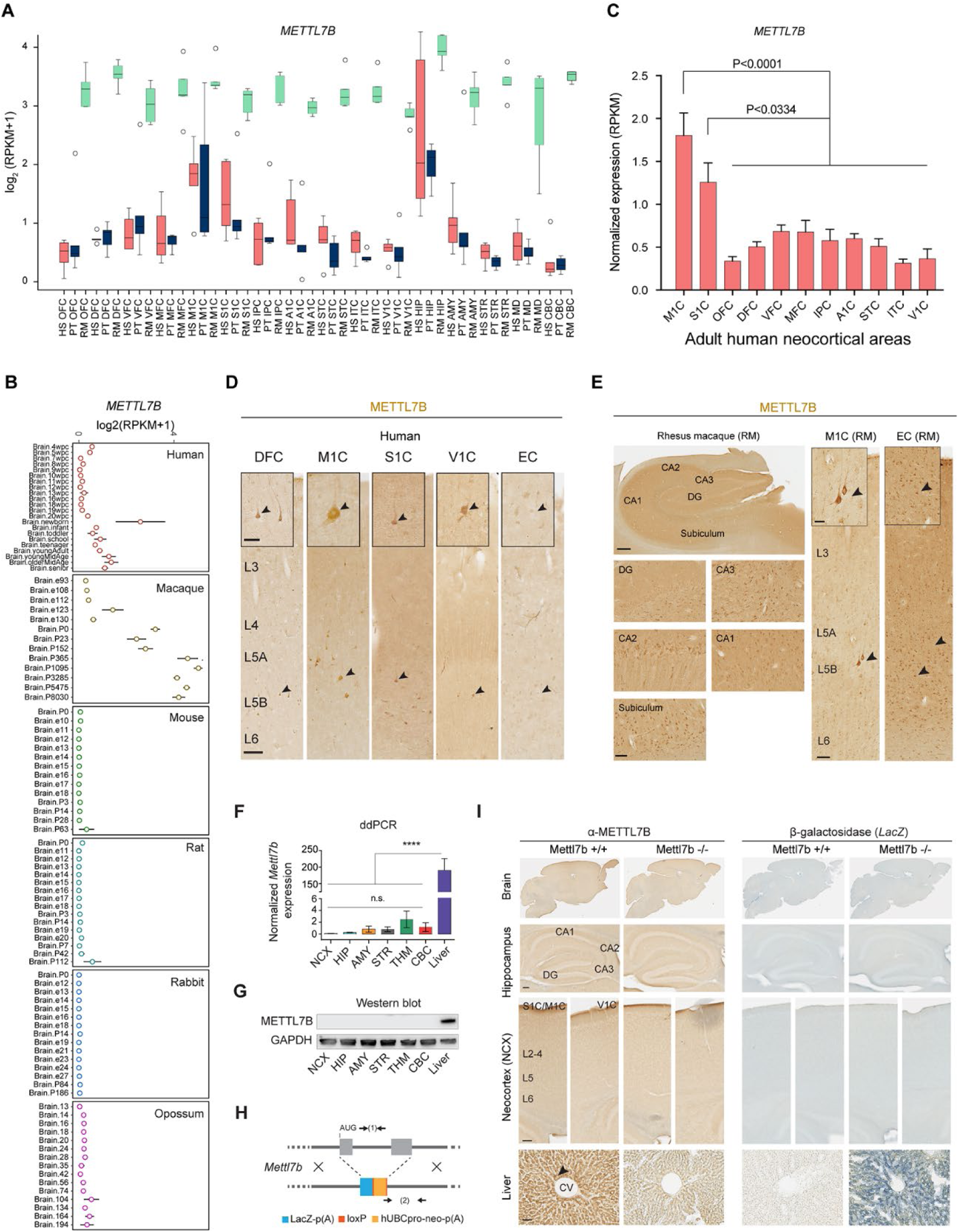
*METTL7B* is expressed in primate brain but not mouse brain. **A**, Exon-array expression of *METTL7B* homologs in human (HS), chimpanzee (PT), and rhesus macaque (RM) brain regions. **B**, RNA-seq expression of *METTL7B* in the brain (forebrain/cerebrum) of multiple species at different developmental stages. Expression data was obtained from Cardoso-Moreira et al. 2019. **C**, RNA-seq expression of *METTL7B* in human NCX. One-way ANOVA with post-hoc Dunnett’s adjustment, all groups N=6, except MFC N=5. Data are means ± SEM. **D**, Prominent immunolabeling of layer 5B (L5B) pyramidal neurons (arrowheads), including Betz and Meynert cells in M1C and V1C, respectively. Scale bars = 150 μm; insets = 50 μm. **E**, METTL7B immunolabeling of hippocampus, Betz cells, and pyramidal neurons in RM brain. Scale bars = 100 μm; inset = 50 μm. **F**, Digital droplet PCR of mouse brain regions and liver, and **G**, immunoblot validation of *Mettl7b* expression in the adult mouse brain and liver showing significantly higher expression in liver with no differences between brain regions. One-way ANOVA with post-hoc Dunnett’s adjustment (****P<0.0001), N=3 per group. All data are mean ± SEM. **H**, Generation of *Mettl7b* knock-out (^-^/^-^) mouse. **I**, Immunostaining reveals Mettl7b protein and lacZ expression in liver. No staining observed in adult mouse brain. Scale bars: brain = 100 μm; liver = 50 μm. CV = central vein. A1C, primary auditory cortex; DFC, dorsolateral prefrontal cortex (aka DLPFC); EC, entorhinal cortex; IPC, posterior inferior parietal cortex; ITC, inferior temporal cortex M1C, primary motor cortex; MFC, medial prefrontal cortex; OFC, orbital prefrontal cortex; S1C, primary somatosensory cortex; STC, superior temporal cortex; V1C, primary visual cortex; VFC, ventrolateral prefrontal cortex.

**Figure S6 (related to Figure 6).**
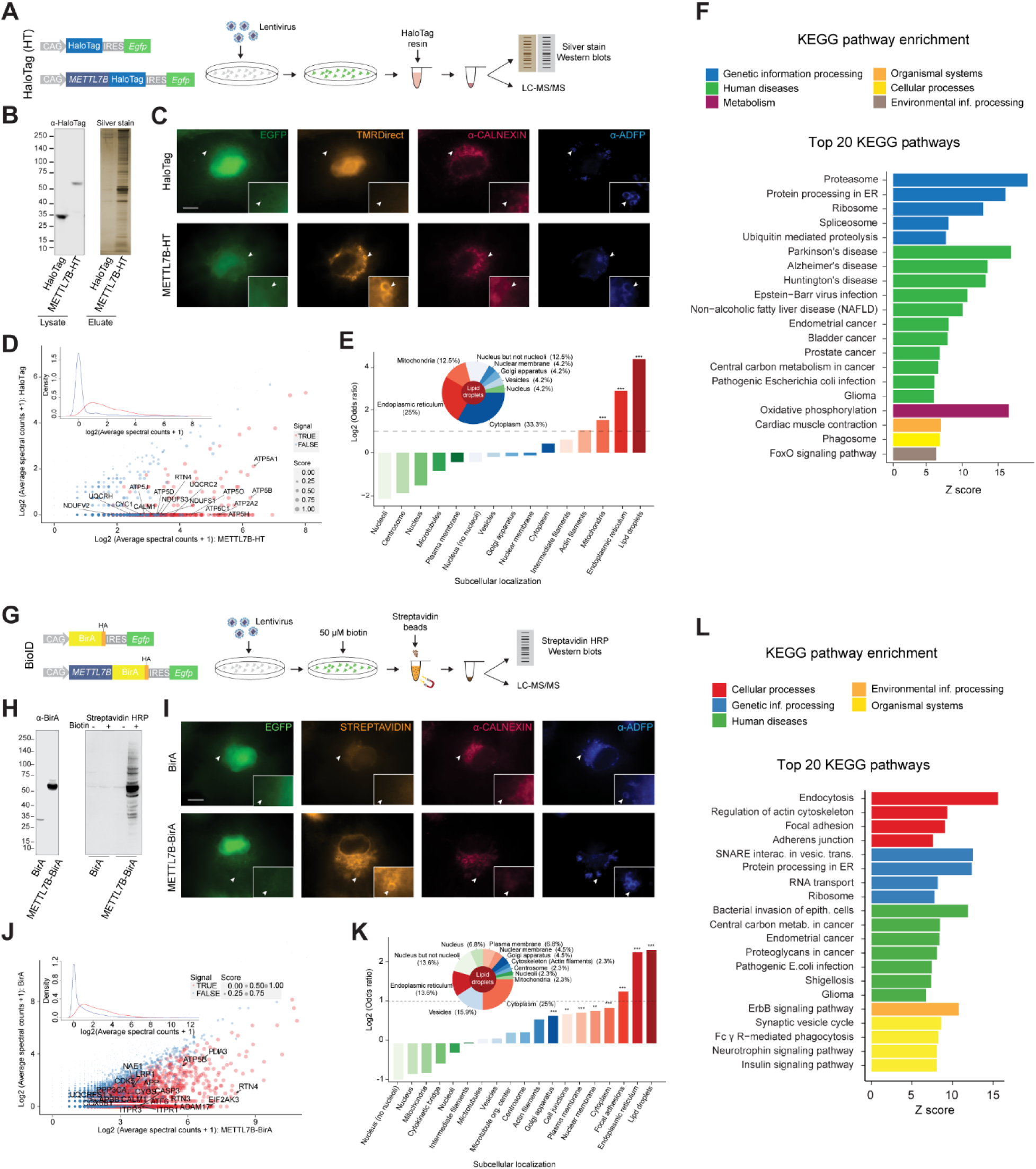
KEGG pathway enrichment of METTL7B interacting proteins. **A,** Schematic of HaloTag (HT) pulldown design. **B,** Immunoblot validation of HT proteins and silver stain of pulldown eluates showing more proteins captured in METTL7B-HT sample. **C,** Immunofluorescence staining showing that METTL7B fusion protein (TMR-Direct) co-localizes with CALNEXIN and ADFP. Scale bars = 10 μm **D,** SAINT analysis distinguishes true METTL7B interactors (red) from false ones (blue) based on MS spectral counts. The figure shows the average spectral counts in 3 test experiments (x axis) and 3 control experiments for all detected proteins. The inset clarifies separation between true METTL7B interactors (red curve) and the false ones (blue curve) in terms of spectral count distribution. **E,** Fold-enrichment test for major subcellular compartments cataloged in Human Protein Atlas database and mammalian cytoplasmic lipid droplet proteomes. The inset shows subcellular composition (%) of LD associated proteins. ***P<0.001. **F,** KEGG pathway enrichment analysis showing molecular pathways involving true interactors are associated with three neurodegenerative diseases: Alzheimer’s, Parkinson’s, and Huntington’s disease. **G**, Schematic of BioID pulldown experimental design. **H**, Immunoblot validation of BioID proteins (α-BirA) and pulldown efficiency (STREPTAVIDIN-HRP) after supplementing cell culture media with 50 μM biotin for 24 hours. **I**, METTL7B-expressing cells exhibit vast biotinylation of endogenous proteins (STREPTAVIDIN) which co-localize with CALNEXIN and ADFP, ER and LD markers, respectively. Scale bars = 10 μm **J**, SAINT analysis distinguishes true METTL7B interactors (red) from false ones (blue) based on MS spectral counts. The figure shows the average spectral counts in 3 test experiments (x axis) and 3 control experiments for all detected proteins. The inset clarifies separation between true METTL7B interactors (red curve) and the false ones (blue curve) in terms of spectral count distribution. **K**, Fold-enrichment test for major subcellular compartments cataloged in Human Protein Atlas database and mammalian cytoplasmic lipid droplet proteomes. The inset shows subcellular composition (%) of LD associated proteins. ***P<0.001, **P<0.01. **L)** KEGG pathway enrichment analysis.

**Figure S7 (related to Figure 6).**
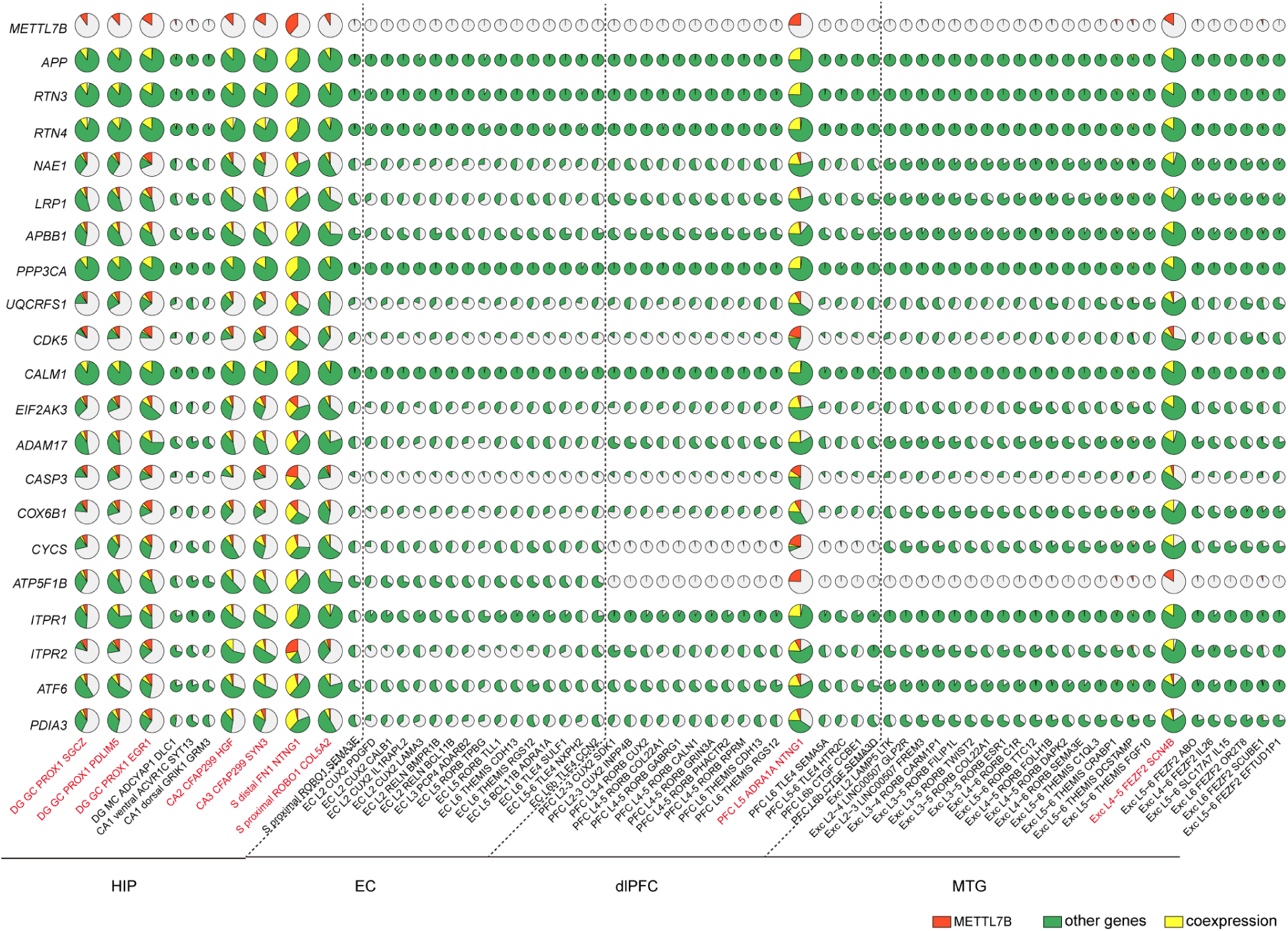
Co-expression of *METTL7B* and its interactors in single cells. Pie charts showing the percent of cells expressing *METTL7B* but not certain *METTL7B* interactor (red), the percent of cells expressing each of the *METTL7B* interactors but not *METTL7B* (green), as well as the percent of cells co-expressing *METTL7B* and certain interactor (yellow) out of all cells within the subtypes of HIP, EC, MTG and PFC. Each row represents a gene and each column denotes a subtype.

## SUPPLEMENTAL TABLES

**Table S1 (related to Figure 4) | Tissue samples used in adult neurogenesis analyses.**

**Table S2 (related to Figure 5) | Complete list of genes exhibiting brain region-specific up- or down-regulated expression.**

**Table S3 (related to Figure 6) | LC-MS/MS data from BioID and HaloTag experiments.**

**Table S4 (related to Figure 6) | SAINT analysis of LC-MS/MS data from BioID and HaloTag experiments.**

**Table S5 (related to STAR methods) | Primer and probe sequences.**

## Methods

### Human, non-human primate tissue

Human samples were obtained from the collections of the Sestan and Rakic laboratories and from Javier DeFelipe’s collection in the Instituto Cajal in Madrid (Spain). Non-human primate brain specimens were obtained from the tissue collection of the Sestan laboratory. All clinical histories, tissue specimens, and histological sections were evaluated to assess for signs of disease, injury, and gross anatomical and histological alterations.

Frozen archival tissue human specimens were used for snRNA-seq. No obvious signs of neuropathological alterations were observed in any of the specimens considered and analyzed in this study. For all other specimens, regions of interest were sampled from frozen tissue slabs or whole specimens stored at −80 °C. To ensure consistency between specimens, all dissections were performed by the same person. Frozen tissue slabs were kept on a chilled aluminum plate during dissections. EC and four hippocampal subregions (DG, CA 2-4, CA1, and Sub) were microdissected as previously reported (Kang et al., 2011)from fresh frozen post-mortem human brains previously cut into 1-cm thick serial, coronal sections, and snap frozen in isopentane (J. T. Baker).

Fresh tissue specimens for histology were fixed with 4% paraformaldehyde/PBS followed by 30% sucrose/PBS. No obvious signs of neuropathological alterations were observed in any of the human, or macaque specimens analyzed in this study. The postmortem interval (PMI) was defined as hours between time of death and time when tissue samples were fresh frozen or started to undergo fixation process.

All human (*Homo sapiens*) brain specimens (18 Y female, 23 Y male, 23 Y male, 42 Y male, 42 Y male, 50 Y female, 51 Y male, 63 Y male, 69 Y female, 74 Y male, 78 Y female, 79 Y female, 79 Y female, and one case where sex and age are unknown; PMIs up to 24 hours) were de-identified and collected from clinically unremarkable donors and one case that died in status epilepticus. Tissue was collected following the guidelines provided by the Yale Human Investigation Committee (HIC) for the Sestan and Rakic collection or by the European Union for DeFelipe’s samples from Spain. Tissue was collected and handled in accordance with ethical guidelines and regulations for the research use of human brain tissue set forth by the NIH (http://bioethics.od.nih.gov/humantissue.html) and the WMA Declaration of Helsinki (http://www.wma.net/en/30publications/10policies/b3/index.html). Appropriate informed consent was obtained and all available non-identifying information was recorded for each specimen.

All studies using non-human primates were carried out in accordance with a protocol approved by Yale University’s Committee on Animal Research and NIH guidelines. The postmortem interval (PMI) was defined as hours between time of death and time when tissue samples were frozen or started to undergo fixation process. Rhesus macaque (*Macaca mulatta*) brain samples were collected postmortem from 2 adult specimens (4 Y male, 8 Y female; PMIs up to 2 hours). Macaque brains were post-fixed by immersion in 4% paraformaldehyde/PBS for approximately 24 hours, followed by 30% sucrose/PBS, and stored at +4°C.

### Anatomical definition of sampled subregions of the hippocampal formation and entorhinal **cortex**

***<u>The dentate gyrus (DG)</u>*** was sampled from the posterior part of the anterior third of the hippocampal formation. It included the suprapyramidal blade, infrapyramidal blade and crest of as well as all three layers: molecular, granular, and polymorphic. The polymorphic layer contained only the superficial part, while the deeper part of the hilus of the DG was dissected as part of the proximal portion (nearer DG) of the CA2-4 region.

***<u>Cornu Ammonis (CA) 2-4 region</u>*** was sampled after DG was dissected and contained the remaining part of the hippocampal proper anlage until approximately CA1 region, including all three layers: molecular, pyramidal and stratum oriens. The proximal portion, corresponding to CA4, contained most the tissue of the hilus of the DG.

***<u>CA1 region</u>*** (Sommer’s sector) was sampled from approximately the border of CA2 to the subiculum, comprising the most distal (further away from the DG) portion of cornu Ammonis. The border between CA1 and CA2 is difficult to reliably identify and thus small pieces of the neighboring CA2 and, vice versa, could have been occasionally present in the sample.

***<u>The subiculum (S)</u>*** forms the most distal part of the hippocampal formation that is adjacent to CA1 region. It was sampled adjacent to the CA1 sample and contained all cortical layers and the superficial region of underlying white matter.

***<u>The entorhinal cortex (EC)</u>*** spreads over both the gyrus ambiens and a considerable part of the parahippocampal gyrus. The EC samples were collected from the middle portion of the parahippocampal gyrus of the same tissue slab used to dissect the subregions of the hippocampal formation, corresponding to the proper entorhinal subregion and Brodmann area 28. The EC was also defined by presence of numerous wart-like elevations (verrucae hippocampi) on the surface of the gyrus. Samples contained all cortical layers and the superficial region of underlying white matter.

### Brain cell nuclei isolation

The brain cell nuclei were isolated according to our previous protocol (Li et al., 2018; Zhu et al., 2018) with some modifications. Hippocampal regions (DG, CA1, CA2-4, Subiculum) and adjacent entorhinal cortex were dissected from three frozen adult human brains (table S). In order to avoid experimental bias and evenly dissociate the tissue for cell nuclei isolation, whole tissue was finely pulverized to powder in liquid nitrogen with mortar and pestle (Coorstek #60316, #60317). All buffers were ice cold and all reagents used for consequent nuclear isolation were molecular biology grade unless stated otherwise. 5 - 10 mg of pulverized tissue was added into 5 ml of ice-cold lysis buffer consisting of 320 mM sucrose (Sigma #S0389), 5 mM CaCl2 (Sigma #21115), 3 mM Mg(Ace)2 (Sigma #63052), 10mM Tris-HCl (pH 8) (AmericanBio #AB14043), protease inhibitors w/o EDTA (Roche #11836170001), 0.1 mM EDTA (AmericanBio #AB00502), RNAse inhibitor (80U/ml) (Roche #03335402001), 1mM DTT (Sigma #43186), and 0.1% TX-100 (v/v) (Sigma#T8787). DTT, RNAse Protector, protease inhibitors, and TX-100 were added immediately before use. The suspension was transferred to Dounce tissue grinder (15ml volume, Wheaton #357544; autoclaved, RNAse free, ice-cold) and homogenized with loose and tight pestles, 30 cycles each, with constant pressure and without introduction of air. The homogenate was strained through 40 um tube top cell strainer (Corning #352340) which was pre-wetted with 1ml wash buffer: (250 mM sucrose (Sigma #S0389), 25 mM KCl (Sigma #60142), 5mM MgCl2 (Sigma #M1028), 20mM Tris-HCl (pH 7.5) (AmericanBio #AB14043; Sigma #T2413), protease inhibitors w/o EDTA (Roche #11836170001), RNAse inhibitor (80U/ml) (Roche #03335402001), 1mM DTT (Sigma #43186)). Additional 4 ml of wash buffer was added to wash the strainer. Final 10 ml of solution was mixed with 10 ml of 50% Optiprep (Axis-Shield# 1114542) solution (50% iodixanol (v/v), 250 mM sucrose (Sigma #S0389), 25 mM KCl (Sigma #60142), 5mM MgCl2 (Sigma #M1028), 20mM Tris-HCl (pH 7.5) (AmericanBio #AB14043; Sigma #T2413), protease inhibitors w/o EDTA (Roche #11836170001), RNAse inhibitor (80U/ml) (Roche #03335402001), 1mM DTT (Sigma #43186)) by inverting the tube 10x and carefully pipetted into 2 centrifuge tubes (Corning #430791). The tubes were centrifuged at 1000g, for 30 min at 4°C on centrifuge (Eppendorf #5804R) and rotor (Eppendorf #S-4-72). Upon end of centrifugation, the supernatant was carefully and completely removed and total of 5 ml of resuspension buffer (250 mM sucrose (Sigma #S0389), 25 mM KCl (Sigma #60142), 5mM MgCl2 (Sigma #M1028), 20mM Tris-HCl (pH 7.5) (AmericanBio #AB14043; Sigma #T2413), protease inhibitors w/o EDTA (Roche #11836170001), RNAse inhibitor (80U/ml) (Roche #03335402001), 1mM DTT (Sigma #43186)) was added carefully on the pellets in tubes and centrifuged at 1000g, for 10 min at 4 °C on the same centrifuge and rotor. Supernatants were then carefully and completely removed, pellets were gently dissolved by adding 100 ul of resuspension buffer (see above) and pipetting 30x with 1ml pipette tip, pooled and filtered through 35 um tube top cell strainer (Corning #352340). Finally, nuclei were counted on hemocytometer and diluted to 1 million/ml with sample-run buffer: 0.1% BSA (Gemini Bio-Products #700-106P), RNAse inhibitor (80U/ml) (Roche#03335402001), 1mM DTT (Sigma #43186) in DPBS (Gibco #14190).

### Single nucleus microfluidic capture and cDNA synthesis

The nuclei samples were placed on ice and taken to Yale Center for Genome Analysis core facility and processed within 15 minutes for single nucleus RNA sequencing with targeted nuclei recovery of 10000 nuclei, respectively, on microfluidic Chromium System (10x Genomics) by following the manufacturer’s protocol (10x Genomics, CG000183_Rev_A), with Chromium Single Cell 3’ GEM, Library & Gel Bead Kit v3, (10x Genomics #PN-1000075) and Chromium Single Cell B Chip Kit (10x Genomics #PN-1000074), Chromium i7 Multiplex Kit (10x Genomics #PN-120262) on Chromium Controller (10x Genomics). Due to limitations imposed by source RNA quantity, cDNA from nuclei was amplified for 14 cycles.

### Single nucleus RNA-seq library preparation

Post cDNA amplification cleanup and construction of sample-indexed libraries and their amplification followed manufacturer’s directions (10x Genomics, CG000183_Rev_A), with the amplification step directly dependent on the quantity of input cDNA.

### Sequencing of libraries

In order to reach sequencing depth of 20000 raw reads per nucleus, single nucleus libraries were run using paired end sequencing with single indexing on the HiSeq 4000 platform (Illumina) by following manufacturer’s instructions (Illumina; 10x Genomics, CG000183_Rev_A). To avoid lane bias, multiple uniquely indexed samples were mixed and distributed over several lanes. RNA-seq data is deposited at http://psychencode.org and NCBI dbGAP Accession phs000755.v2.p1.

### Quantification and Statistical Analysis

#### Single nuclei expression quantification and quality control

We quantified the expression levels of genes in each potential nucleus represented by a cellular barcode using the 10X Genomics CellRanger pipeline (version 3.0.2). Reads were mapped to human reference genome GRCh38 (Ensembl release 98) and quantified in units of Unique Molecular Identifiers (UMIs) based on the combined exon-intron human annotation. We took advantage of the enhanced cell-calling methodology in CellRanger to distinguish true cells from damaged or empty droplets. Specifically, RNA content distribution of each barcode was compared to the background concentration which was generalized from extremely low RNA-containing barcodes, and was subsequently classified as damaged if comparable profiles were seen. To further rule out low-quality cells, we excluded nuclei with mitochondrial content greater than 10%. This loose criterion was set as we aimed to incorporate certain cell types into analyses such as endothelial cells which were shown to be prone to high mitochondrial content (Velmeshev et al., 2019). Additional filtering procedure was performed after clustering and low-dimensional embedding (see below) to eliminate small cell clusters collectively displaying elevated mitochondrial and ribosomal gene expression and showing no signals of reasonable cell types.

### Normalization, dimensionality reduction and clustering

We normalized the raw UMI counts into log-transformed Transcripts Per Million (TPM) using the ‘NormalizeData’ function in the R package Seurat (scaling factor equals to 10,000) (version 3.1.0) (Butler et al., 2018). To position all nuclei in a two-dimensional representation reflecting their transcriptomic similarities (Fig. 1B-1D), the top 2,000 highly variable genes were obtained by the Seurat function ‘FindVariableFeatures’ with the default variance stabilizing process for each of the three human individuals. We further integrated nuclei from the three humans on the basis of their anchor features summarized from each individual via the function ‘IntegrateData’ and embedded ensuing nuclei in the Uniform Manifold Approximation and Projection (UMAP) plot using the top 30 principal components (PCs) (‘RunPCA’ function in Seurat followed by the function ‘RunUMAP’). To cluster nuclei according to their nearest transcriptomic neighbors, we searched for shared nearest neighbors (SNN) in the PCA space with the neighbor number being 20 and optimized the graph modularity using the Seurat function ‘FindClusters’. In general, we performed an iterative removal-clustering approach to remove nuclei with high mitochondrial or ribosomal contents and without clear cluster-related markers followed by re-clustering of the remaining nuclei. Moreover, cells co-expressing multiple cell-type specific marker genes were manually marked as doublets and excluded from the downstream analytical flow. Lastly, we re-embedded cell types of interest (i.e., excitatory neurons, inhibitory neurons and non-neuronal cells) in the UMAP space and re-clustered them using the same procedure as mentioned above, as this would offer finer details into the cell types we sought to probe into.

### Global across-dataset comparison

We performed global comparisons with the human MTG (Hodge et al., 2019) and dlPFC single nucleus datasets (Li et al., 2018) to investigate the similarities and distinctions among them. We processed the MTG and dlPFC data using the same procedure except that for MTG, the scaling factor during data normalization was set to 1,000,000 to mitigate the bias caused by magnitude differences. For each of the highly variable genes detected in both our data and MTG/dlPFC data, we averaged the TPMs across each subtype within both data and transformed them in the logarithmic space. Spearman correlation coefficients were calculated across these subtypes to further avoid the across-dataset batches and the resulting linkages were exhibited in gradient heat maps (Fig. S1E, S1F).

### Tree construction

To explore the taxonomic relationships among all cell subtypes, we constructed a hierarchical tree by first averaging the gene expression levels across cells of the same subtype. The derived expression was standardized to mean of zero and variance of one within each subtype across the anchor genes selected in the previous integration step. Following this step, we calculated the Euclidean distances between pairwise subtypes, and clustered these subtypes in a structured tree (Fig. 1E) by the ‘hclust’ function in R with the method set to ‘ward.D2’.

### Classification of cell subtypes

We grouped cell clusters with strong signals of *SLC17A7* expression into excitatory neurons. Furthermore, we categorized them into different subtypes through marker gene expression and comparisons with published datasets (Fig. S2A, S2B). Specifically, granule cells were characterized by the predominant composition of dentate gyrus nuclei and prominent expression of *PROX1*. Mossy cells were described by the principal origin from dentate gyrus and exclusive expression of *ADCYAP1*. We subsequently classified granule cells into three subtypes characterized by the high expression of *SGCZ*, *PDLIM5* and *EGR1*, respectively. Excitatory neurons from CA fields were arranged mainly according to subfields: CA3 pyramidal neurons (co-expression of *CFAP299* and *SYN3*), CA2 pyramidal neurons (co-expression of *CFAP299* and *HGF*), dorsal CA1 pyramidal neurons (co-expression of *GRIK1* and *GRM3*), and ventral CA1 pyramidal neurons (co-expression of *ACVR1C* and *SYT13*). For the subiculum excitatory neurons, we categorized them into three subtypes: one distal (away from CA1) (*FN1*+) subtype and two proximal ones (*ROBO1*+). Of note, the spatial registrations of CA and subiculum cell subtypes were achieved on the basis of previous transcriptomic studies of hippocampal pyramidal neurons (Cembrowski et al., 2016a; Cembrowski et al., 2016b; Cembrowski et al., 2018). With regards to entorhinal excitatory neurons, we classified them by two means. First, we aligned them with excitatory neurons from single nucleus data of human MTG using the same procedure as described above. Second, we examined the subtype-specific marker genes in both our excitatory neurons and related literature reports. Specifically, two layer 2 subtypes were classified as *RELN*+ and one as *CALB1*+ (Witter et al., 2017). Other upper-layer subtypes were depicted based on marker gene expression of *LAMA3*, *PDGFD*, *IL1RAPL2*, and *PCP4* (Ramsden et al., 2015; Tang et al., 2015; Ohara et al., 2018). The middle-to-deep layer subtypes were delineated by the specific gene expression of *RORB*, *THEMIS*, *ADRA1A*, and *TLE4*.

Cell clusters showing high GAD1 expression were then assigned as inhibitory neurons. Inhibitory neuron clusters were first classified to major groups based on the expression of three canonical function markers (*PVALB*, *SST*, *VIP*) as well as *LAMP5*, a marker mostly representing a group of neurogliaform inhibitory neurons and recently being adopted as a major inhibitory neuron marker (Tasic et al., 2018; Hodge et al., 2019). For a cluster expressing two markers simultaneously (eg. InN LAMP5 NMBR cluster expresses both SST and LAMP5), it was assigned to the same major group of the neighboring cluster in the hierarchical tree. Additionally, we used LHX6 (a medial ganglionic eminence marker) and NR2F2 (a caudal ganglionic eminence marker) to classify the rest of the inhibitory neuron clusters which do not express these markers. We also identified an inhibitory neuron cluster with no evident markers and showing high mitochondria counts, indicative of low-quality cells, which accordingly was termed as “InN unknown”. Finally, each inhibitory neuron cluster was named after the combination of major group marker (eg. SST, VIP) and one top exclusive cluster marker (eg. ANO2). Apart from these inhibitory neuron clusters, we also identified a neuron cluster co-clustered with inhibitory neurons showing strong signals of *RELN*, *NDNF,* highly indicative of Cajal Retzius cells.

The remaining nuclei were collectively referred to as non-neuronal cells. We classified these nuclei into four big groups based on marker gene expression of *SOX10* (oligodendrocyte lineage-related cells), *AQP4* (astrocytes), *PTPRC* (immune cells) and *RGS5* (endothelial cells) (Fig. S2E, S2F). The first group was further subdivided by the expression of *PDGFRA* (oligodendrocyte precursor cells, OPCs), *GPR17* (committed oligodendrocyte precursor cells, COPs), and *MOBP* (oligodendrocytes), as in Fig. S2H. We additionally grouped OPCs and oligodendrocytes into specific subtypes according to the high expression of specific genes: *EGR1* and *GRIA4* for OPCs; *CPXM2*, *SLC5A11*, *LINC01098* and *LAMA2* for oligodendrocytes. For astrocyte subtype specification, we classified them by the laminar distribution: *GFAP*+ ones located in deep layers and *CHRDL1*+ ones in upper layers (Fig. S2G) (Lanjakornsiripan et al., 2018). Regarding immune cells, we used marker genes *C1QB*, *F13A1*, *LYZ* and *SKAP1* to deconstruct them into microglia, macrophages, myeloid cells and T cells, respectively (Fig. S2I). Microglia were further subdivided via specific gene expression of *P2RY12* and *CD83*. In terms of vasculature lineage, we employed combinational expression of genes to sort them into arterial endothelial cells (*DKK2*+), endothelial cells (*CLDN5*+ and *VWF*+), pericytes (*CLDN5*+ and *ABCC9*+), venous smooth muscle cells (*ABCC9*+ and *P2RY14*+), arterial smooth muscle cells (*ACTA2*+ and *TAGLN*+) and vascular and leptomeningeal cells (*COL1A2*+ and *COL1A1*+) (Fig. S2J) (Vanlandewijck et al., 2018).

### Cell subtype comparisons among HIP, EC, MTG and dlPFC

To explore the transcriptomic divergence across HIP, EC, MTG and dlPFC for all cell subtypes, we constructed a network demonstrating the relationships among the subtypes in the four brain regions based on the extent of overlap of their specific marker genes. In detail, in each region we first determined the marker genes of each subtype using the ‘FindAllMarkers’ function in Seurat. Subsequently, we generated a similarity matrix representing the overlap between marker genes of pairwise subtypes across all regions, followed by the visualization of this matrix in the form of a network via the R package ‘igraph’ through the force-directed graphopt algorithm (Fig. 2A-F). Especially, for excitatory neuron types we displayed their connections in a between-region manner (HIP and EC, EC and MTG, and MTG and dlPFC). To further examine the cell subtype connections across different regions, in each brain region we focused on marker genes detected in at least one subtype and assessed their expression across all subtypes of remaining brain regions visualized in heat maps (Fig. S3A). Additionally, given the upper- and deep-layer marker genes identified in MTG, we calculated the percentages of genes in each subtype of each region where expression was greater than the expression constraint of 75% quantile across all expression values. Furthermore, we evaluated the expression of marker genes from intratelencephalic/intracerebral (IT) neurons and non-IT neurons of MTG in all subtypes of the four regions through first averaging the expression of each gene across cells of the same subtype and then displaying the median values across IT markers/Non-IT markers in scatter plots (Fig. S3C).

### Exclusive markers of cluster InN SST ADAMTS12

To find hippocampus-specific transcriptome features in the cluster InN *SST ADAMTS12*, we first sought to confirm the enrichment of this cluster in hippocampus by integrating inhibitory neurons from across the hippocampal-entorhinal system with those from MTG and dlPFC using the ‘fastMNN’ function from the batchelor R package (Fig. 3C) (Haghverdi et al., 2018). Following this confirmation, we identified a set of markers exclusively expressed in this cluster as compared to other interneuron clusters in hippocampus and SST-expressing interneuron clusters in MTG or dlPFC. To do so, we first calculated a specificity score for each gene in each cluster to assess the specificity of gene expression in each cluster (Li et al., 2018). For cluster InN *SST ADAMTS12*, we then selected those genes with a specificity score in the 99% quantile and that showed a 1.5 fold or greater difference between this specificity score and the maximum specificity score of that gene in any other cluster. To further refine this list of genes exhibiting highly selective expression in In *SST ADAMTS12*, we next removed from this list those genes that were also highly specific (maximum specificity score in the 99^th^ quantile) for any population of *SST*-expressing neurons in MTG or dlPFC. The expression of these markers in each cell was then generalized as a single AUROC score calculated via AUCell package (Aibar et al., 2017).

### Expression of DCX and proliferation markers in HIP and EC

To check the expression of *DCX* in the cell types of HIP and EC, we interrogated *DCX* expression in all excitatory or inhibitory neurons and next in all DG or EC cell types, all of which were re-embedded in the UMAP space (Fig. 4A-D). During visualization, the expression threshold (i.e., log-transformed TPM) was set to one to highlight the apparent expression of *DCX* in corresponding cell types. Quantitatively, we evaluated the percentage of DG inhibitory neurons or GC expressing *DCX* under different thresholds in units of UMIs (1, 2 and 3). We further examined the cells expressing *NES*, *MKI67* or *DCX*, the three markers labeling neural stem cells (NSC), neural progenitor cells (NPC), and migrating and immature granule cells (IM) respectively, in DG and EC. These cells were uncovered in each subtype of DG or EC, and the marker genes over the granule cell maturation were checked specifically for those cells (Fig. 4E, 4F **and** SFB, S4C).

To test whether DCX-expressing cells showed enriched expression of *NEUROD1, NEUROD2, SOX11, CALB2, FXYD7, STC1, NCAM1 and PROX1*, we compared, in each cluster, the proportion of *DCX*-expressing and *DCX*-negative cells also expressing each of those markers by means of a Fisher Exact’s test (Fig. S4A). P-values were adjusted using Bonferroni correction.

### Analysis of bulk tissue transcriptomic datasets

Gene expression analysis was performed on an exon array (Kang et al., 2011) and PsychENCODE RNA-seq datasets (Li et al., 2018). Gene expression values from exon array were used to rank protein-coding genes based on a region-specific upward or downward temporal trajectory. Time periods 3-15 were collapsed into three time groups: prenatal (periods 3-7), early postnatal (periods 8-12), and adult (periods 13-15). We used limma (Smyth, 2005) to run a regression that included the time group and brain region, as well as the region-group interactions, as factors. To select genes with an increase in expression across time groups in a single region, genes were ranked by their region-group interaction coefficient. We filtered to genes for which the time group coefficient was above an arbitrary cutoff of −0.05 to remove cases where the high region-group interaction simply offsets an overall negative decrease in expression across the time groups. We also filtered out genes where more than one region-group interaction coefficient was above 0.01 to restrict the ranking to increases in expression that are unique to a single brain region. To identify genes where expression decreases across time groups in a region-specific manner, similar criteria were used, reversing the sign of the cutoff values and the direction of the comparisons. Gene expression values from BrainSpan RNA-seq dataset were used to compare *METTL7B* expression within multiple neocortical regions during adulthood (time periods 13-15). Statistical analysis was performed using one-way ANOVA.

### Generation of knockout mice and tissue processing

All experiments with mice were performed in accordance with a protocol approved by Yale University’s Committee on Animal Research. Targeted embryonic stem (ES) cells (Mettl7b^tm1(KOMP)Vlcg^) were obtained from Knockout Mouse Project (KOMP) repository. Chimeric mice were generated by blastocyst injection of ES cells at Yale Genome Editing Center (YGEC). Mice were bred for germline transmission to generate gene knockout mice. Genotyping was performed using the TUF/TUR primer set (145 bp) for the wild-type allele and the NeoFwd/SD primer set (351 bp) for the *Mettl7b* deletion allele.

Both wild type and *Mettl7b* mutant mice were reared in group housing in a 12h light:12h dark cycle and provided food and water ad libitum with veterinary care provided by Yale Animal Resource Center. Only mice in good, healthy condition, as approved by Yale Animal Resource Center, were used for breeding and experimentation. Multiple breeding pairs were maintained and siblings were never mated to increase genetic diversity, and prevent unintended selection for features that could affect results. Both sexes were used and randomly assigned for all experiments. Adult mice were anesthetized and intracardially perfused with ice-cold PBS and 4% PFA. All mouse brain tissue specimens were fixed by immersion in 4% PFA overnight at 4 °C and sectioned at 50 μm using a vibratome (Leica).

### *In situ* hybridization

Human brain tissue samples were fixed in 4% PFA overnight at 4 °C and sectioned at 30 μm using a Leica VT1000 S vibratome. The RNA probes complementary to human *METTL7B* cDNA (NM_152637.2) were labeled with digoxigenin-UTP (Roche). After acetylation, sections were hybridized with the probes at 63 °C for 16 hours. Following hybridization, the riboprobes were immunolabeled with anti-digoxigenin-AP conjugate and the signal was developed with NBT/BCIP overnight in dark.

### Immunolabeling and histology

For METTL7B immunohistochemistry, tissue sections were processed using ImmPRES Excel Amplified HRP Polymer Staining Kit (Anti-Rabbit IgG, MP-7601-15, Vector Laboratories) per manufacturer’s protocol. For mouse α-β-galactosidase (*lacZ*) stain, tissue sections were blocked with blocking solution (5% normal donkey serum, 1% BSA, 0.1% glycine, 0.1% lysine, and 0.3% Triton X-100 in PBS) for 1 hour and incubated with primary antibodies and biotinylated secondary antibodies. The signal was amplified with Vectastain ABC-AP kit (AK-5000, Vector Labs) and developed with Vector Blue AP kit (SL-5300, Vector Labs) per manufacturer’s protocol. DCX and GAD1 immunohistochemistry was performed with anti-DCX antibodies raised in guinea pig (EMD Millipore AB2253; 1:1000) and anti-GAD1 antibody raised in goat (R&D AF2086; 1:200) in 3% normal donkey serum, 0.25% Triton X-100 in PBS). Antigen retrieval (20 mins in citrate buffer pH 6 at 95C) was required for optimal results. Antibody detection was achieved with biotinylated secondary antibody and Streptavidin conjugated (Jackson Immunoresearch) for DCX and anti-goat secondary antibodies (Jackson Immunoresearch) for GAD1. DAPI was used for nuclear staining. All histology samples were imaged on Aperio ScanScope system or imaged on a Zeiss LSM 510 confocal microscope. Cell culture samples were fixed with ice-cold 4% paraformaldehyde (PFA) for 10 minutes at room temperature, blocked for 30 minutes at RT with blocking solution (5% normal donkey serum, 1% BSA, 0.1% glycine, 0.1% lysine, and 0.3% saponin in PBS), incubated with primary and appropriate Alexa Flour-conjugated secondary antibodies, and imaged on Zeiss LSM 510 confocal microscope.

### Plasmids

For expression of *METTL7B*, full length cDNA (NM_152637.2) was inserted into pCAGIG (a gift from Connie Cepko, Addgene #11159) (Matsuda and Cepko, 2004). For lentiviral generation, pFUGW (a gift from David Baltimore, Addgene #14883) (Lois et al., 2002) was digested with PacI, 3’ overhangs removed with Klenow (NEB) to form blunt ends, and additionally digested with BsrGI to release hUBC promoter and EGFP. The CAG-IRES-EGFP was removed from pCAGIG and ligated into pFUGW. For protein pulldown experiments, BirA-HA and HaloTag constructs were PCR-amplified from pcDNA3.1-MCS-BirA(R118G)-HA (a gift from Kyle Roux, Addgene #36047) (Roux et al., 2012) and pHTC-CMVneo-HaloTag (G7711, Promega), respectively, and ligated into pFUGW-CAG. For overexpression of wild type APP, full length APP (NM_201414.2) and mCherry cDNA were PCR amplified and ligated into pFUGW-CAG in place of IRES-EGFP. For doxycycline inducible expression of METTL7B, cDNA fragments [Ampicilin resistance, high-copy-number Origin of replication, SV40 poly(A), and IRES-EGFP from pCAGIG; rTetR and tight TRE promoter from pCW57.1 (a gift from David Root, Addgene #41393); hPKG promoter (M60581.1); METTL7B (NM_152637.2); bGH poly(A) from pFUGW] were PCR amplified, ligated, and circularized (pDTET-METTL7B).

### Lentiviral purification and generation of stable cell lines

Ten 15-cm dishes of sub-confluent Lenti-X 293T cells (Clontech) were used for each purification. pFUGW-CAG specific plasmids (BirA, METTL7B-BirA, HaloTag, METTL7B-HaloTag) along with pMD2.G, pRSVrev and pMDLg/pRRE (a gift from Didier Trono, Addgene #12259, #12253, #12251) (Dull et al., 1998) were transfected at 1:1:1:1 molar ratio using PolyJet (SignaGen). Cell culture media containing lentiviral particles (LVP) was collected at 48- and 60-hours post-transfection and filtered through 0.2 μm filter to remove cellular debris. Filtered supernatants were centrifuged at 100,000g for 2 hours. One milliliter of PBS was laid over LVP pellet and left overnight at 4 °C. Next day, resuspended pellets were centrifuged through 30% sucrose gradient to further purify the virus. Lentiviral titers were determined by transducing Lenti-X 293T cells and calculating titer from FACS data between 1-10% infection rate using formula: Titer (IU/ml) = (# cells seeded x dilution factor x % GFP-positive cells) / (volume of virus solution added).

For pulldown experiments, 50,000 ReNcell CX (EMD Millipore) cells were plated on a laminin coated 24-well plate in triplicate wells. Cells were transduced with lentiviral particles at MOI of 10 in a 150 μL of cell culture media supplemented with 10 μg/mL of protamine sulfate (#02194729, MP Biomedicals) and saved as ReN-CAG-BirA, ReN-CAG-METTL7B-BirA, ReN-CAG-HaloTag, and ReN-CAG-METTL7B-HaloTag stable cell lines.

For *APP* overexpression, N2a cells were transduced at MOI of 40. After propagation, eight million cells were sorted at Yale Flow Cytometry Facility, Yale University, on a BD SORP FACSAria 2 cell sorter (Special Order Research Product) using the 100-μm nozzle and a sheath pressure of 20 p.s.i. BD FACSDiva Software was used to acquire and analyze samples. Gates were set to remove cell doublets and particle debris. Cell selection was based on mCherry signal intensity and top one percent expressing cells were saved as the N2a-APP cell line.

### Affinity capture of proteins

For BioID and HaloTag experiments, two million cells (ReN-CAG-BirA, ReN-CAG-METTL7B-Bira, ReN-CAG-HaloTag, ReN-CAG-METTL7B-HaloTag) were plated on four laminin coated 10-cm dishes. BioID pulldown was performed per protocol (Roux et al., 2013). At near confluency, cell culture media was supplemented with 50 μM biotin (B4639, Sigma-Aldrich). The next day, cells were rinsed twice with PBS, detached with Accutase (Millipore) for 10 minutes at 37 °C, centrifuged at 200 g for 3 minutes, rinsed with PBS, and centrifuged again. Bead-protein conjugates were resuspended in 50 mM ammonium bicarbonate. HaloTag pulldown was performed per manufacturer’s protocol (G6500, Promega). Proteins were eluted by resuspending HaloTag resin in 50 μL of 8 M urea prepared in 50 mM ammonium bicarbonate and shaking for 30 minutes at room temperature. Ten percent fractions of BioID and HaloTag eluates were saved for immunoblot and silver stain analysis.

### Mass spectrometry and proteomic data analysis

BioID and HaloTag tryptic digestion was performed using the optimized method from the original published method(Kim et al., 2014). Proteins were reduced by adding 2 μl of 0.5M Tris(2-carboxyethyl)phosphine (TCEP) at 30 °C for 60 min. The reaction was cooled to room temperature (RT) and proteins were alkylated in the dark for 30 min by adding 4 μl of 0.5M Iodoacetamide. Sample volume was adjusted by adding 350 μl of 50 mM Ammonium Bicarbonate to dilute the 8M urea to 1M before trypsin digestion. Mass spectrometry grade trypsin (Promega) was added for overnight digestion at 30°C using Eppendorf Thermomixer at 700 rpm. Formic acid was added to the peptide solution (to 2%), followed by desalting by C18 TopTip (TT10C18.96, PolyLC) and finally dried on a SpeedVac. Tryptic peptides were resuspended in 100 μl of 2% Acetonitrile in 0.1% formic acid. Ten microliters of total tryptic peptides were used in triplicate runs for the 1D LC-MS/MS analysis, consisting of an EASY-nLC 1000 HPLC Acclaim PepMap peptide trap with a 25 cm-2μm Easy-Spray C18 column, Easy Spray Source, and a Q Exactive Plus mass spectrometer (all from Thermo Fisher Scientific). A 230-min gradient consisting of 5–16%B (100% acetonitrile) in 140 min, 16-28% in 70 min, 28-38% in 10 min, 38-85% in 10 min was used to separate the peptides. The total LC time was 250 min. The Q Exactive Plus was set to scan precursors at 70,000 resolution followed by data-dependent MS/MS at 17,500 resolution of the top 12 precursors.

#### Protein Identification and data analysis

The LC-MS/MS raw data of two technical replicates was combined and submitted to Sorcerer Enterprise v.3.5 release (Sage-N Research Inc.) with SEQUEST algorithm as the search program for peptide/protein identification. SEQUEST was set up to search the target-decoy UniProt Human Reviewed (v. March 2015) protein fasta database using trypsin for the enzyme and with the allowance of up to 2 missed cleavages, semi tryptic search, fixed modification of 57 Da for cysteines to account for carboxyamidomethylation and precursor mass tolerance of 50 ppm. Differential search included 226 Da on lysine for biotinylation (BioID samples), 16 Da for methionine oxidation, and 14, 28 and 42 Da on lysine for mono-, di- and tri-methylayion. The search results were viewed, sorted, filtered, and statically analyzed by using comprehensive proteomics data analysis software, Peptide/Protein prophet v.4.02 (ISB). The minimum trans-proteomic pipeline (TPP) probability score for proteins was set to 0.9 to assure very low error (less than FDR 2%) with good sensitivity. The differential spectral count analysis was done by QTools, an open source SBP in-house developed tool for automated differential peptide/protein spectral count analysis(Brill et al., 2009) and the protein prophet peptide report was utilized to report biotinylated peptides. The LC-MS/MS raw data were also submitted to Integrated Proteomics Pipelines (IP2) Version IP2 v.3 (Integrated Proteomics Applications, Inc.) with ProLucid algorithm as the search program (Xu et al., 2006) for peptide/protein identification. ProLucid search parameters were set up to search the UniProt Human Reviewed (v. March 2015) protein fasta database including reversed protein sequences using trypsin for enzyme with the allowance of up to 2 missed cleavages, semi tryptic search, fixed modification of 57 Da for cysteines to account for carboxyamidomethylation and precursor mass tolerance of 50 ppm. Differential search included 226 Da on lysine for biotinylation (for BioID samples), 16 Da for methionine oxidation, and 14, 28 and 42 Da on lysine for mono-, di- and tri-methylayion. The search results were viewed, sorted, filtered, and statically analyzed by using DTASelect for proteins to have protein FDR rate of less than 2.5% (Tabb et al., 2002). Differential label-free proteomics data analysis was done by IP2-Census, Protein Identification STAT COMPARE (Park et al., 2008) using two technical replicates. This result was a label-free quantification analysis, of duplicate technical data for each sample; using spectral count analysis with t-test and Gene Ontology analysis (Robinson et al., 2004).

#### Identification of true pulldown proteins based on mass spectrometry spectral counting data

We discriminated true prey-bait interactions from false interactions in the Halotag and BioID pulldowns by using Significance Analysis of INTeractome (SAINT) method (Choi et al., 2011; Teo et al., 2014). Briefly, the SAINT method utilizes MS/MS spectral counting data and models true and false prey-bait interactions as separate Poisson distributions to obtain the probability of a *true* protein-protein interaction based on Bayesian statistical inference. The estimated probability provides a quantitative measure of the confidence of prey-bait interactions such that false interactions can be filtered out in a statistically-controlled manner. Upon applying the SAINT method to MS/MS spectral count data available from each pulldown experiment system, we identified 275 (out of 3 cases and 3 controls) and 1795 (3 cases and 3 controls) proteins as true METTL7B interactors from Halotag and BioID pulldowns, respectively, at Bayesian False Discovery Rate (BFDR) of 5%.

#### Subcellular localization analysis

To characterize subcellular localization of the true METTL7B interactors, we performed fold-enrichment test for major subcellular compartments cataloged in the Human Protein Atlas database (Uhlen et al., 2015) and mammalian lipid droplet proteomes (Hodges and Wu, 2010). Human Protein Atlas provides genome-wide analysis of major subcellular localization information of human proteins based on immunofluorescent stained cells. It consists of 20 main subcellular compartments and 10,003 proteins (www.proteinatlas.org). To make the fold-enrichment test comparable across Human Protein Atlas and the mammalian lipid droplet proteome datasets, we merged the mammalian lipid droplet protein list to Human Protein Atlas dataset as a separate subcellular localization category and used the entire Human Protein Atlas subcellular localization records uniformly as a null (background) set. We found that 73.8% (203/275) and 77.7% (1384/1795) of true METTL7B interactors from HaloTag and BioID pulldown experiments had matching HGNC gene symbols in Human Protein Atlas. Of the 152 mammalian cytoplasmic lipid proteins(Hodges and Wu, 2010), 80 proteins had matching HGNC gene symbols in the Human Protein Atlas. Twenty-three (HaloTag) and 37 (BioID) true METTL7B interactors were identified to be among 80 lipid droplet proteins in the Human Protein Atlas database.

#### Validation of pulldown experiment using a publicly available protein-protein interaction database (comparison with the BioGRID protein-protein interaction database)

We evaluated the performance of SAINT method by benchmarking the true METTL7B interactors against non-redundant physical BioGRID protein-protein interaction network (Stark et al., 2006). We computed the significance of interactions between proteins from the true METTL7B interactor set and the rest of the proteins (background set) in the protein-protein interaction (PPI) network by using binomial proportions test Z-score as follows (Abul-Husn et al., 2009):

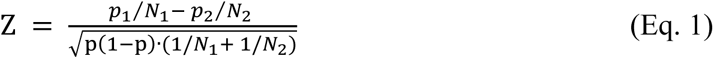

where

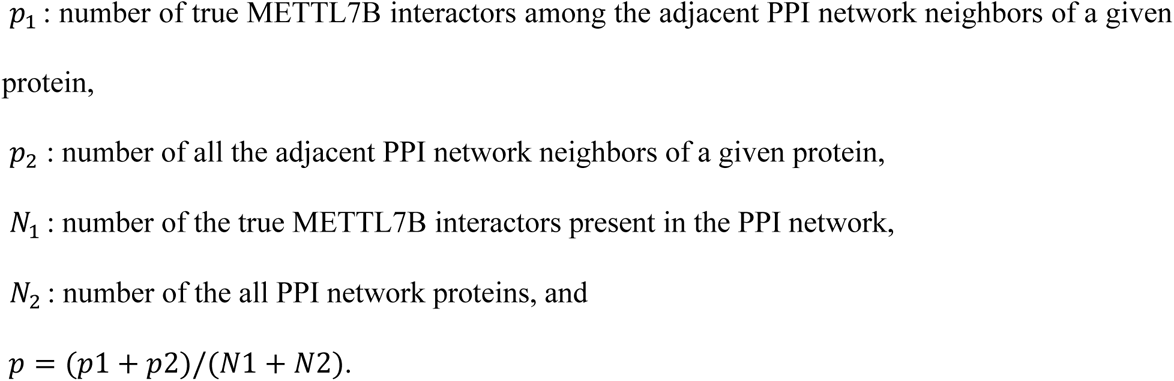

The Z-score thus provides an approximate quantitative measure of how significantly a given protein in the PPI network interacts with the true METTL7B interactors in the immediate neighborhood of the protein-protein interaction network compared to the background proteins in the protein-protein interaction network. We found that the true METTL7B interactors tend to interact much more significantly to each other than to the rest of proteins in the protein-protein interaction network (Wilcoxon rank sum test p-value < 2e-16, data not shown). This indicates that the true METTL7B interactors are significantly clustered and proximal to each other in the protein-protein interaction network as expected.

#### KEGG pathway enrichment analysis

To gain an insight of molecular processes associated with the true METTL7B interactors, we performed KEGG pathway enrichment analysis using the binomial proportion test Z-scores as weights of proteins in a given pathway. The rationale for using such a weight is that the proteins interacting significantly more with true METTL7B interactors play proportionally important role in specifying the biological context represented by the true METTL7B interactors. To this end, we assigned biological context specificity scores to all KEGG pathways in a similar manner used in (Ideker et al., 2002) as follows:

First, we assigned a pathway-level Z score (*Z_A_*) as a sum of all individual KEGG pathway protein member Z scores divided by the square root of number of member proteins (*k*)

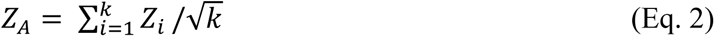

Second, each pathway-level Z score (*Z_A_*) was calibrated against null expected mean (*μ_k_*) and standard deviation (*σ_k_*) for a given pathway consisting of *k* member proteins, which are empirically estimated from 10,000 randomly selected gene sets of size *k* from the BioGRID protein-protein interaction network.

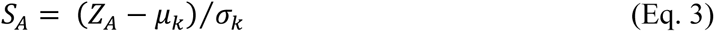

The analysis results are then organized by KEGG BRITE functional classification scheme. This allows to compare the relative significance of KEGG pathways (Kanehisa and Goto, 2000; Kanehisa et al., 2016; Kanehisa et al., 2017) within functionally related BRITE categories as well as between distinct BRITE categories.

#### Spatial clustering analysis

We examined spatial clustering between METTL7B true pulldown proteins and KEGG pathways in protein-protein interaction network and evaluated the significance of the spatial clustering between them as a function of network distance. To this end, we extended existing implementation of spatial statics K*^net^* function (Cornish and Markowetz, 2014) to allow us to examine spatial correlation of two types of points (e.g., a set of proteins in a KEGG pathway and a set of METTL7B true pulldown proteins) in the protein-protein interaction network. We define spatial correlation of two groups (e.g., group *a* and group *b*) of points as follows,

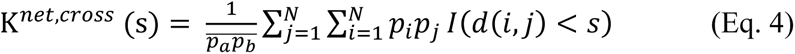

Here, *p_i_* and *p_j_* denote group membership for all the vertex of protein-protein interaction network. (e.g., *p_i_* =1 and *p_j_* =1 if a protein is a member of group *a* and group *b*. *p_i_* =0 and *p_j_* = 0 otherwise). The *d*(*i, j*) denotes network distance between a pair of proteins in two groups, *a* and *b*. *I*(*d*(*i*, *j*) < *s*) is an indicator function with value of 1 for a pair of proteins in the two groups closer than network distance s and 0 otherwise. 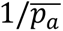 and 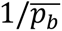 are normalization constants giving a weight of each protein in two groups 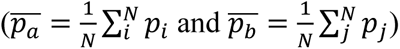. To estimate the statistical significance of K*^net, cross^*(*s*) across network distance, we randomly permuted group membership in the network and obtained the significance as a Z-score: 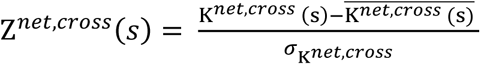 where 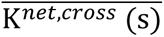 and *σ_k_^net, cross^* are mean and variance of empirical null expectation obtained from the random permutations. Thus, Z*^net, cross^* (*s*) provides an insight about a profile of spatial overlap between the two groups of molecular signatures in protein-protein interaction network.

### ELISA

To determine the differences in Aβ generation, 20 000 N2a-APP cells per well were plated in 96-well plate. After four hours, cells were transfected with pDTET-EGFP or pDTET-METTL7B-EGFP using PolyJet reagent (SignaGen). Four hours post transfection, cells were rinsed twice with PBS and incubated in fresh media supplemented with 200 nM doxycycline. After 48 hours, conditioned media was collected and supplemented with protease inhibitor cocktail (P-2714, Sigma-Aldrich). Cells were lysed in Cell Extraction Buffer (Thermo Fisher Scientific) supplemented with protease inhibitor cocktail. Cell culture supernatants were analyzed in duplicates on Ab40 and Ab42 colorimetric ELISAs per manufacturer’s protocol (KHB3481 and KHB3544, Invitrogen). Aβ concentrations were normalized per total cell protein concentrations, measured by the Rapid Gold BCA Protein Assay (#A53225, Pierce).

### Immunoblotting and silver stain

#### Tissue sample preparation

Tissue was lysed in PBS with 0.01% Tween-20 and protease inhibitor cocktail (P-2714, Sigma-Aldrich), and sonicated in two sessions (30 pulses at an output level of 3 using a Microson Ultrasonic Cell Disruptor [Misonix]) with 1-minute rest on ice between sessions. Samples were centrifuged at 14 000 g for 10 minutes at 4 °C. Total protein concentrations were measured by the Bradford assay (#23246, Pierce).

#### Immunoblotting

Samples were mixed with NuPAGE LDS Loading Buffer (NP0007) supplemented with 50 mM DTT, incubated at 72 °C for 10 minutes, and loaded on 4-12% Bis-Tris gel (NP0321, Thermo Fisher Scientific). Proteins were transferred to a 0.2 μm PVDF membrane (#162-0218, Bio-Rad), blocked with 5% non-fat milk or BSA in 1% TBST buffer, and blotted with appropriate primary and secondary HRP-conjugated antibodies. The signal was developed with SuperSignal West Pico Plus Chemiluminescent Substrate (#34577, Pierce) and visualized on G:BOX Chemi XRQ (Syngene) system.

#### Silver stain

5% of HaloTag eluates were prepared as above and electrophoresed on 4-12% Bis-Tris gel. Gel was processed using Silver Stain for Mass Spectrometry kit per manufacturer’s instructions (#24600, Pierce).

#### SAM assay

Custom made recombinant METTL7B was expressed in *E. Coli* ArcticExpress and purified from inclusion bodies by GenScript. Recombinant RTN3, RTN4, LRP1, and APP peptide were purchased directly from vendors. SAMfluoro Methyltransferase Assay (786-431, G-Biosciences) was performed per manufacturer’s instructions using ∼2 μg of METTL7B and ∼1 μg of substrate protein. Recombinant proteins were incubated with or without METTL7B in triplicate wells. Assay was performed at 37 °C and resorufin fluorescence was measured on GloMax Multi Detection System (Promega) plate reader with an excitation wavelength of 530-540 nm and an emission wavelength of 585-595 nm.

#### RNA isolation and digital droplet PCR

Total RNA was extracted from human and mouse brain tissue samples, or cultured cells, using RNAeasy Plus Mini Kit (#74134, Qiagen) per manufacturer’s protocol. RNA concentrations and quality were determined using R6K ScreenTape (#5067-5576, Agilent) and TapeStation analyzer (Agilent). cDNA was synthesized from 1 μg of total RNA using SuperScript III First-Strand Synthesis kit (#18080051, Invitrogen) and random primers. Digital droplet PCR was performed using QX200 Droplet Digital PCR (Bio-Rad) and data was normalized to *TBP* expression. PCR amplification was performed using primer sets and probes listed in Table S4.

